# Crowder-induced Conformational Ensemble Shift in *Escherichia Coli* Prolyl-tRNA Synthetase

**DOI:** 10.1101/603548

**Authors:** L. M. Adams, R. J. Andrews, Q. H. Hu, H. L. Schmit, S. Hati, S. Bhattacharyya

## Abstract

The effect of macromolecular crowding on the structure and function of *Escherichia coli* prolyl-tRNA synthetase (Ec ProRS) has been investigated using a combined experimental and theoretical method. Ec ProRS is a multi-domain enzyme; coupled-domain dynamics is essential for efficient catalysis. To gain an insight into the mechanistic detail of the crowding effect, kinetic studies were conducted with varying concentrations and sizes of crowders. In parallel, spectroscopic and quantum chemical studies were employed to probe the “soft-interactions” between crowders and protein side chains. Finally, the dynamics of the dimeric protein was examined in the presence of crowders using a long-duration (70 ns) classical molecular dynamic simulations. The results of the simulations revealed a significant shift in the conformational ensemble, which is consistent with the “soft-interactions” model of the crowding effect and explained the observed alteration in kinetic parameters. Collectively, the present study demonstrated that the effects of molecular crowding on both conformational dynamics and catalytic function, are correlated. This is the first report where molecular crowding has been found to impact the conformational ensemble in the multi-domain Ec ProRS, a member of aminoacyl-tRNA synthetase family, which is central to protein synthesis in all living cells. The present study affirmed that the effect of crowders should be considered while investigating the structure-dynamics-function relationship in modular enzymes.

## INTRODUCTION

AARSs are multi-domain enzymes that play a critical role in protein biosynthesis in all living organisms; they catalyze the covalent attachment of amino acids to their respective tRNAs (1, 2). A member of AARSs is prolyl-tRNA synthetase (ProRS), which catalyzes the ligation of proline to tRNA^Pro^ in a two-step reaction. The first step of the reaction is the formation of the enzyme-bound prolyl-adenylate complex (Pro-AMP) and pyrophosphate (PP_i_) in the presence of adenosine triphosphate (ATP) (eq. 1). In the second step, the activated amino acid is transferred to the 3’-end of tRNA^Pro^ (eq. 2) forming aminoacylated tRNA (Pro-tRNA^Pro^) (1, 2).

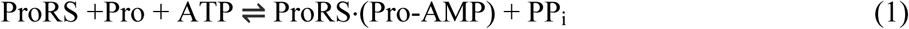

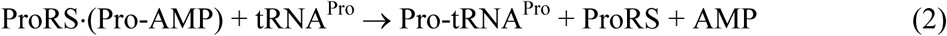

Many AARSs including ProRSs undergo a large-scale conformational change upon substrate binding (3–5). Previous studies from our research group have demonstrated that the dynamic coupling between distant domains is crucial for effective catalysis (6, 7). However, most experimental and theoretical studies are generally carried out in aqueous environments that are completely different from what is found in-vivo. The interior of a cell is crowded with many biological macromolecules and metabolites. The cytoplasm of cells such as *Escherichia coli* have a macromolecule concentration of up to 300-400 g/L, and that around 20-30% of the intracellular volume is occupied by macromolecules (8, 9). Recent studies revealed that crowded environment have noticeable effect on protein stability (10–12), folding (13), binding affinity (14, 15), and catalysis (13, 16–18). Specifically, the size, structure, concentration, and chemical nature of crowding agents contribute to changes in stability, folding, conformational equilibrium, and catalysis when compared to dilute conditions (19–22). A study by Ferreira et al. suggested that the chemical nature and concentration of the crowding agents induce changes in solvent dielectric properties, which in turn modulates “soft-interactions” between biological macromolecules and crowding agents (23).

The above findings pose new questions about the role of macromolecular crowding on the structure and function of Ec ProRS. There are two main questions that emerge: is the coupled-dynamics impacted by crowding and if impacted, does it alter the protein’s function? To shed light onto these questions and to gain molecular-level understanding of the effects of macromolecular crowding on Ec ProRS function, an investigatory approach with three major components − enzyme kinetics, intrinsic tryptophan fluorescence spectroscopy, and molecular dynamic (MD) simulations have been employed. Kinetic assays were performed to explore the impact of size and concentration of crowding agents on the catalytic efficiency of Ec ProRS. In parallel, interactions between protein side chains and crowding agents were studied through intrinsic tryptophan fluorescence measurements and validated through approximate quantum chemical studies. Finally, the conformational dynamics in presence and absence of crowding agents were studied by long-duration molecular dynamics simulation. The ensembles of evolved conformational populations were examined, and their impact on enzyme functions was assessed. The present study has provided a comprehensive assessment of the impact of molecular crowding on the conformation, dynamics, and catalytic function of Ec ProRS.

## MATERIALS AND METHODS

All crowding agents were purchased from Sigma Aldrich, except for polyethylene glycol (PEG) 8000 (Fisher Scientific). Proline (≥ 99%) was also from Sigma Aldrich. Both [γ- ^32^P] ATP and [^32^P] PP_i_ were purchased through Perkin Elmer, Shelton, CT.

### Overexpression and purification of Ec ProRS

Wild-type (WT) Ec ProRS was overexpressed in SG13009 (pREP4) competent cells using 0.1 mM isopropyl β-D-thiogalactoside for 4 hours at 37 °C. Histidine-tagged WT Ec ProRS was purified using Talon cobalt affinity resin column; 100 mM imidazole was used to elute the protein (24, 25). The Bio-Rad protein assay (Bio-Rad Laboratories) was used to determine total concentration of protein. An active-site titration was performed to determine the concentration of active protein (26).

### Enzyme kinetics

#### Crowding agent concentration variation

The ATP-PP*_i_* exchange assay was performed, following the protocol described elsewhere, to examine the effect of increasing concentrations of crowding agents on proline activation (eq.1) by Ec ProRS (27). In ATP-PP_i_ exchange assay, radiolabeled PP_i_ (^32^PP_i_) and non-radiolabeled ATP were used and the percent product (Pro-AMP) formation at 20 minutes post-initiation of the reaction was measured in the presence of crowding agents. In the present study, the amount of Pro-AMP formed was indirectly measured by monitoring the amount of ^32^P-ATP formed *via* the reverse reaction of eq. 1 (ProRS + Pro + ATP ⇌ ProRS**·**(Pro-AMP) + PP_i_). The percent product formation was calculated from the ratio of the Pro-AMP formation in presence of the crowding agent and in absence (i.e. the regular buffer solution). Reactions containing 0.75 mM proline, 10 nM WT Ec ProRS, and the crowding agent were incubated at 37 °C for 20 minutes before being quenched with 0.4 M PPi, 15% HClO4, and 3% activated charcoal. A range of 100-300 g/L dextrose, sucrose and Ficoll were used, while PEG 8000 was present in a range of 50-200 g/L. The same concentrations of proline and enzyme were used in dilute condition as well. The reaction mixture containing a given crowding agent and substrates (proline and ATP) in the absence of enzyme was considered as a control. For all crowding agents, the stability of the enzyme was examined by incubating the enzyme with a given crowding agent for 30 min and analyzing for any degradation using polyacrylamide gel electrophoresis (data not shown).

#### Proline concentration variation to determine the kinetic parameters for proline activation

ATP-PP*_i_* exchange assays, were performed following the method described previously.(27) The impact of the crowding agent on proline activation (eq. 1) efficiency was ascertained by comparing the *K*_M_ and *V*_max_ of WT Ec ProRS relative to that in the absence of any crowding agent (i.e. the dilute solution). These assays employed proline concentrations between 0.125 mM −1.00 mM, 10 nM of WT Ec ProRS, and 200 g/L of crowding agents (dextrose, sucrose, and Ficoll 70). Concentrations higher than 50 g/L for PEG 8000 resulted in a dramatic reduction in catalytic activity, preventing the consistent and accurate determination of kinetic parameters. Therefore, the final concentration was maintained at 50 g/L for PEG 8000. Reaction mixtures were incubated at 37 °C for 4, 8, 12, 16, and 20 minutes before being quenched with 0.4 M PP_i_, 15% HClO_4_, and 3% activated charcoal (27). Kinetic parameters were calculated from Lineweaver-Burk plots. All kinetic assays were performed either in duplicate or triplicate (when the difference between the two measured parameters was greater than 10%). A Student’s t-test with two-tailed statistical analysis was performed on the data from both the proline variation study and the crowding agent concentration variation study.

#### Polyethylene glycol size variation

To investigate the impact of the size of crowding agent, without changing the chemical nature of crowding agents, a size variation study was conducted that utilized PEG 600, 1500, 4000, 6000, 8000, and 20,000. Again, the ATP-PP*_i_* exchange assay was performed (27), where constant concentrations of 0.75 mM proline, 10 nM WT Ec ProRS, and 100 g/L PEG were maintained. Reactions were incubated at 37 °C for 20 minutes before being quenched with 0.4 M PP_i_, 15% HClO_4_, and 3% activated charcoal (27). The quantity of Pro-AMP (nmol) formed was indirectly measured by monitoring the amount of ^32^P-ATP formed and compared to the *R*_h_ (Å) of each PEG (Supplementary Table 1) (28). Reactions with dilute conditions were performed as a reference.

### Intrinsic tryptophan fluorescence spectroscopy

Intrinsic tryptophan fluorescence was measured using a PerkinElmer LS 55 Fluorimeter (PerkinElmer, Inc., MA, USA). Excitation wavelength was 280 nm, and the emission spectra were collected from 300 nm to 400 nm. Baseline emission spectra (controls) were recorded with solutions that contained all reagents but the WT Ec ProRS enzyme (i.e. crowders, buffer, and salt) to eliminate the effect of impurities. Fluorescence solutions contained crowding agents of concentrations ranging from 25-400 g/L, 2 µM WT Ec ProRS, 10 mM phosphate buffer (pH = 7.4), and 100 mM NaCl. Crowding agents from smallest hydrodynamic radius to largest hydrodynamic radius are as follows: dextrose, sucrose, PEG 8000, and Ficoll 70 (Table 1). Experiments were also performed that utilized a combination of two different crowding agents namely, sucrose with Ficoll 70, sucrose with dextrose, and Ficoll 70 with dextrose. Samples were incubated at room temperature for 10 minutes. All experiments were performed in triplicate. The barycentric mean fluorescence wavelength (λ_bcm_) was determined using the equation (eq. 3):

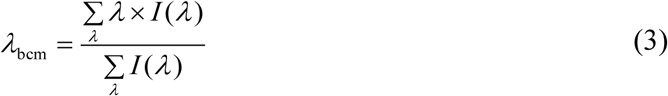

where *λ* is the wavelength and *I*(*λ*) is the emission intensity at that given wavelength. The change in *λ*_bcm_ [Δ*λ*_bcm_ = *λ*_bcm_ (crowder) - *λ*_bcm_ (water)] due to the presence of crowders was examined to monitor changes in the tryptophan’s local environment.

**Table 1.**
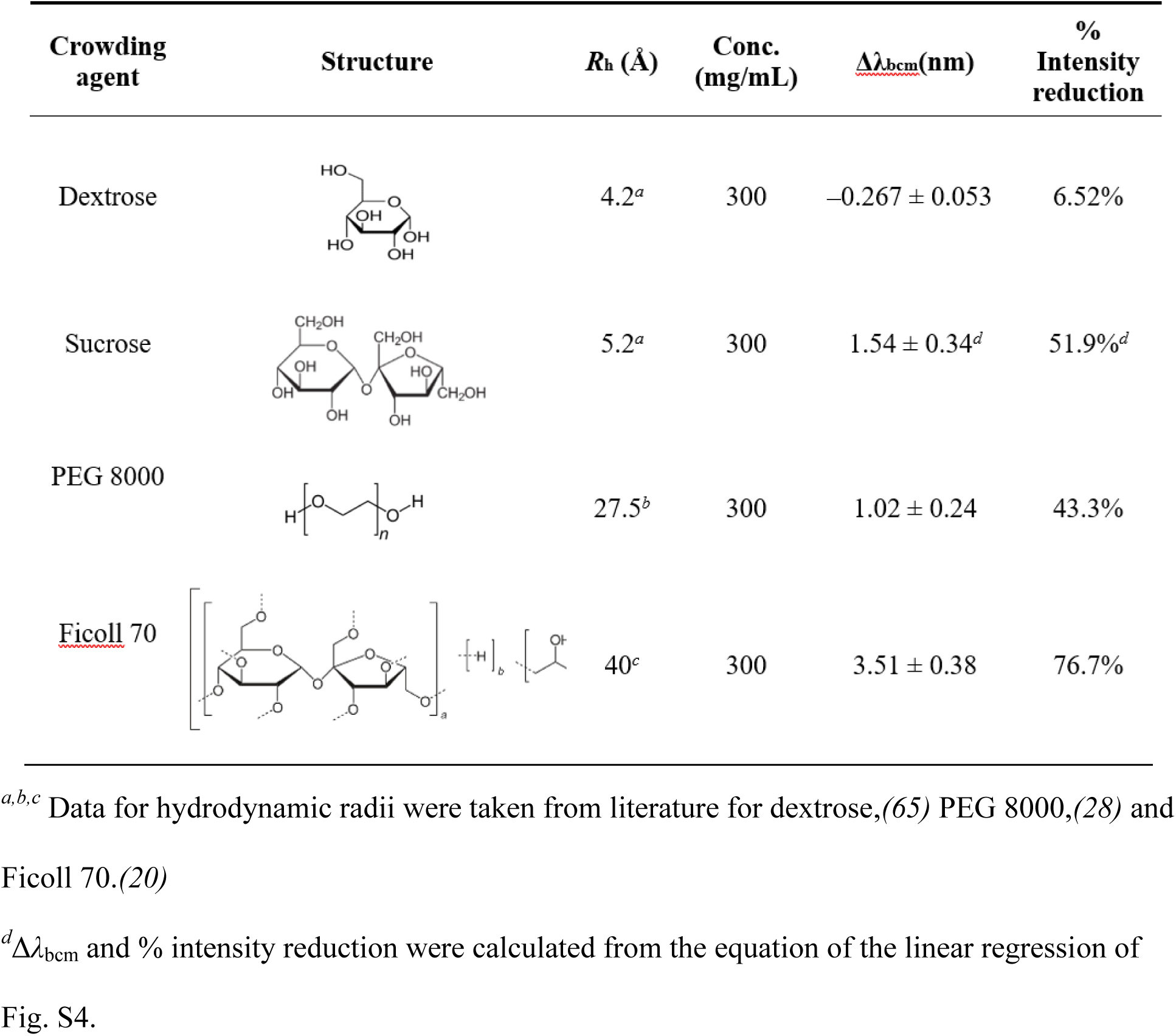
Tryptophan intrinsic fluorescence in the presence of crowding agents.

| Crowding agent | Structure | $R_h$ (Å) | Conc. (mg/mL) | $\Delta\lambda_{bcm}$(nm) | % Intensity reduction |
| --- | --- | --- | --- | --- | --- |
| Dextrose | | 4.2<sup>a</sup> | 300 | $-0.267 \pm 0.053$ | 6.52% |
| Sucrose | | 5.2<sup>a</sup> | 300 | $1.54 \pm 0.34^d$ | 51.9%<sup>d</sup> |
| PEG 8000 | | 27.5<sup>b</sup> | 300 | $1.02 \pm 0.24$ | 43.3% |
| <u>Ficoll 70</u> | | 40<sup>c</sup> | 300 | $3.51 \pm 0.38$ | 76.7% |

<sup>a,b,c</sup> Data for hydrodynamic radii were taken from literature for dextrose,(65) PEG 8000,(28) and Ficoll 70.(20)
<sup>d</sup> $\Delta\lambda_{bcm}$ and % intensity reduction were calculated from the equation of the linear regression of Fig. S4.

### Computational setup

All gas-phase potential energies were calculated using self-consistent density-functional tight-binding theory with dispersion corrections (SCC-DFTB-D) (29, 30). Protein structure was obtained from the protein database (31). Homology models were constructed using the web-based utility SWISS-MODEL (32–34). All structural manipulations were carried out using Visual Molecular Dynamics (VMD) (35). Packmol was used to distribute molecules of sucrose and dextrose randomly throughout the simulation volume (36). Molecular dynamics simulations were carried out using NAMD package using CHARMM36 force field (37–41). Electrostatic energy calculations were carried out using particle mesh Ewald method (42). All solvent accessible surface area calculations were performed with VMD’s dedicated algorithm using a probe size of 1.4 Å. Radial distribution function (RDF) were calculated from the MD trajectory data using VMD. The collective motion was analyzed using principal component analysis of the MD trajectory data using CARMA (43, 44). Backbone root-mean-square-deviation (RMSD) calculations were performed with CARMA using the first frame of the trajectory as the reference.

### Molecular dynamics (MD) simulation

All simulations were performed on a homology model structure of Ec ProRS constructed using Ec ProRS amino acid sequence and *Enterococcus faecalis* ProRS crystal structure (PDB entry: 2J3L) as a template. These simulations involved the dimeric form of the Ec ProRS and were carried out in the presence of two crowding agents, namely, dextrose and sucrose in addition to that in dilute condition. The setup of these systems in the present simulation have been carried out following protocols used in our previous study (7). All structures were explicitly solvated (with TIP3P model) and ionized (with 42 sodium atoms) with VMD plugins, resulting in solvent boxes with dimensions 138 Å × 114 Å × 95 Å for sucrose and 145 Å × 115 Å × 101 Å for dextrose (35). The sucrose and dextrose boxes were built in the same manner, to a final crowding concentration of 200 g/L. Any placements having an atom less than 2.0 Å from any other atom were deleted. After completion of the setup, there were a total of 766 sucrose molecules and 777 dextrose molecules. The total number of water molecules in sucrose and dextrose systems were 31100 and 29205, respectively.

All systems underwent 40 ps of minimization. The 70 ns of MD simulations were run using the NAMD package with time steps of 2.0 femtoseconds. All simulations were run at a constant temperature of 298 K and pressure of 1 bar via NAMD implementation of Langevin dynamics (45) within periodic boundary conditions. The electrostatic calculations involved a grid size of 150 Å × 150 Å × 180 Å. Switching was turned on for electrostatic and van der Waals interactions with a ‘switchdist’ of 10 Å, a cutoff of 14 Å, and a ‘pairlistdist’ of 16 Å.

### Principal component analysis (PCA)

Essential dynamics (46) of the protein was calculated by PCA following procedures as discussed earlier (6). Briefly, a modified trajectory file was prepared from the 70 ns trajectory by removing the overall translational and rotational motions and retaining the information of only C_α_ atoms’ fluctuations. The principal components (or modes) of the motion were obtained by diagonalizing the covariance matrix computed for C_α_ atoms. A clustering of conformations was carried out based on the contribution of the first three principal components and the backbone fluctuations were studied using CARMA (43).

### Potential energy surface calculation

The calculations of potential energy of interactions between tryptophan residues with solvent molecules were carried out in the gas-phase. The interaction potential energy was calculated between a single indole molecule and a water molecule or a crowding agent (dextrose and sucrose). The structural models of these three interacting systems were generated using CHARMM (47) package. Each system was developed by moving the solvent/small molecule from 1 to 12 Å through a direction perpendicular to the plane of the indole by 0.1 Å. A total of 100 single-point potential energy per system were calculated with self-consistent charge-density functional tight-binding with dispersion correction.

## RESULTS AND DISCUSSION

As described earlier, ProRS catalyzes a two-step aminoacylation reaction. The present study has focused on the first step, which involves the formation of Pro-AMP intermediate. This first step of the reaction (eq. 1) is tRNA^Pro^ independent for Ec ProRS (27) and it is the rate-limiting step for class II aminoacyl-tRNA synthetases like ProRS (48). Therefore, we determined the kinetic parameters for the first step of the aminoacylation reaction to explore the impact of crowders on the enzymatic function. This narrowed down the number of complications considerably that may have arisen due to the presence of tRNA^Pro^. With the kinetic assays, we have considered the impact of concentration, size, and the chemical nature of a crowding agent on the Pro-AMP formation i.e. proline activation.

### Decrease in product formation with increasing concentration of crowding agents

There was a significant decrease in product (Pro-AMP) formation in the presence of crowding agents (Fig. 1). General trends show that as the concentration of the crowding agent increases, the product formation decreases (Fig. 1). Polyethylene glycol (PEG) 8000 caused the most significant decrease in product formation (Fig. 1). Even at a lower concentration of 50 g/L, the percentage of product formation was less than that observed with 300 g/L dextrose and sucrose (Fig. 1).

**Figure 1.**
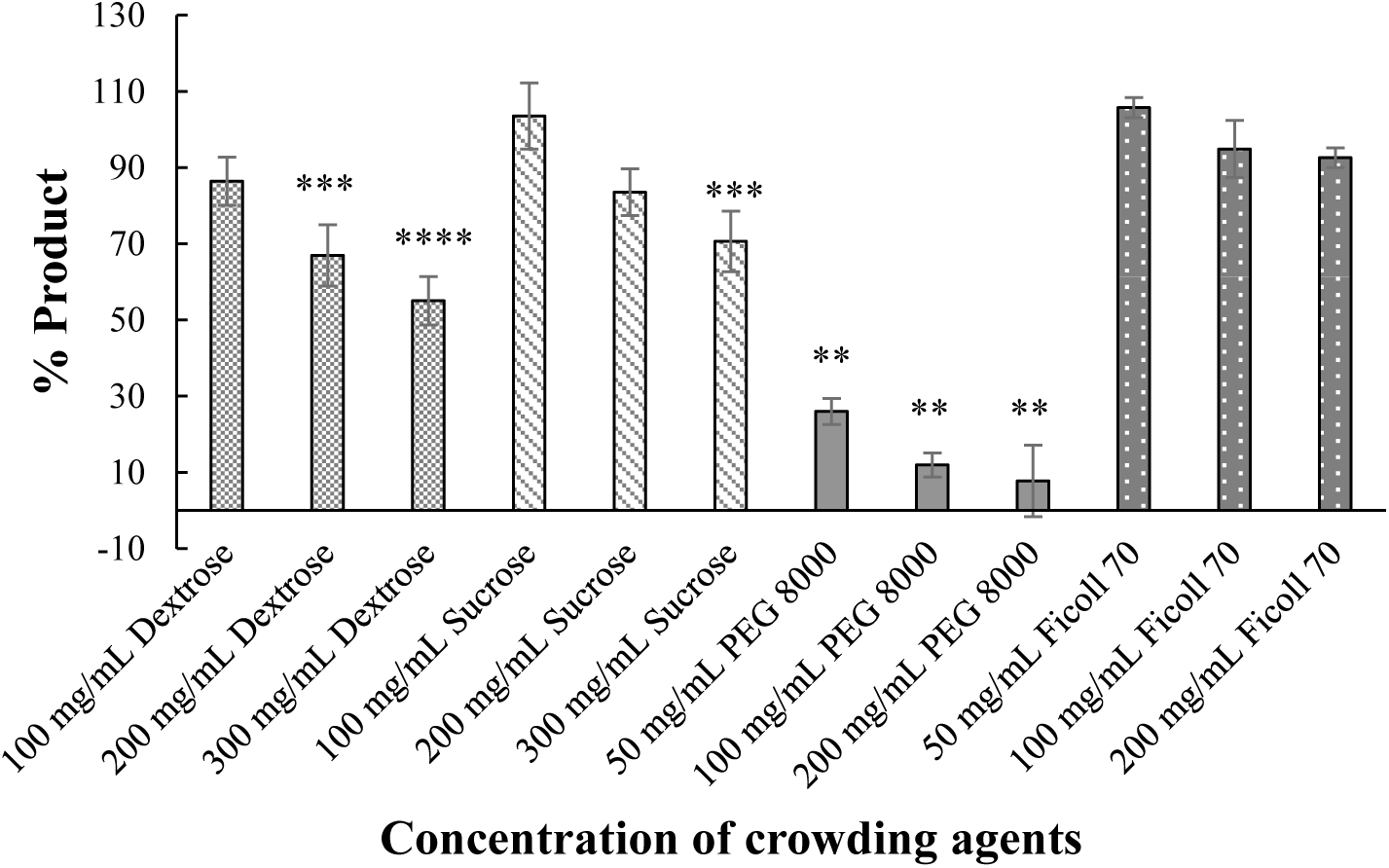
Percent product formation in the presence of variable concentrations of crowding agents. Crowding agent concentration ranged from 100-300 g/L for dextrose, sucrose, and Ficoll 70 and 50-200 g/L for PEG 8000. A concentration of 10 nM WT Ec ProRS and 0.75 mM proline were used. Reactions were incubated at 37 °C and quenched at 20 minutes post-initiation. The percent product formation was calculated by comparing the nmol of Pro-AMP formed after 20 minutes in the presence of crowding agents to that formed in dilute conditions. A two-tailed Student’s *t* test was performed to assess the significance of the difference between the percent product formation in the presence of each individual crowding agent of a specific concentration and the dilute condition. The significance of the results of the *t* test are represented as follows: *p < 0.005, **p < 0.01, ***p < 0.05, ****p < 0.08.

The above results revealed that the product formation decreases with the increasing concentration of crowders, irrespective of their chemical nature or size. One can rationalize this observed decreasing trend by considering the steric effect, which can limit the accessibility to the enzyme active site and thereby impact the product formation. Alternately, the crowding may also influence enzyme conformational equilibrium and internal dynamics. Such variation in conformational equilibrium due to crowding has been reported earlier (21) and could have impacted the product formation by the multi-domain Ec ProRS, where coupled-domain dynamics is known to be important for catalysis (6). In order to have better insight into the impact of crowding agents on prolyl-adenylate formation, the kinetic parameters were determined.

### Impact of crowding on kinetic parameters

Ec ProRS was observed to follow Michaelis-Menten kinetics in the presence of the crowding agents used in the present study. The *K*_M_ and *V*_max_ for proline activation by Ec ProRS were obtained from Lineweaver-Burk plots (Fig. S1 and Table S2). A tighter substrate binding is evident in the presence of crowders as the *K*_M_ was found to be significantly lower (Fig. 2a). The most significant decrease in relative *K*_M_ was obtained with 200 g/L dextrose, resulting in a relative *K*_M_ less than 0.5 (Fig. 2a). The analysis of the relative *K*_M_ revealed that all the four crowding agents lower the *K*_M_ irrespective of crowder size (Fig. 2a). The reaction rate (*V*_max_) of the formation of a prolyl-adenylate decreased for all crowders. For example, in the presence of 50 g/L PEG 8000, the relative *V*_max_ was found to be significantly reduced (> 75%) compared to the dilute condition (Fig. 2b). Also, compared to the dilute condition, the relative *V*_max_ in 200 g/L dextrose and 200 g/L sucrose decreased by about 50% and 25%, respectively. However, the relative *V*_max_ was not significantly different in the presence of 200 g/L Ficoll 70 (Fig. 2b). Compared to the dilute condition, a significant decrease in *k*_cat_/*K*_M_ was observed in the presence of 50 g/L PEG 8000 whereas, an overall increased catalytic efficiency of Ec ProRS was noted in the presence of dextrose, sucrose, and Ficoll 70 (Fig. 2c).

**Figure 2.**
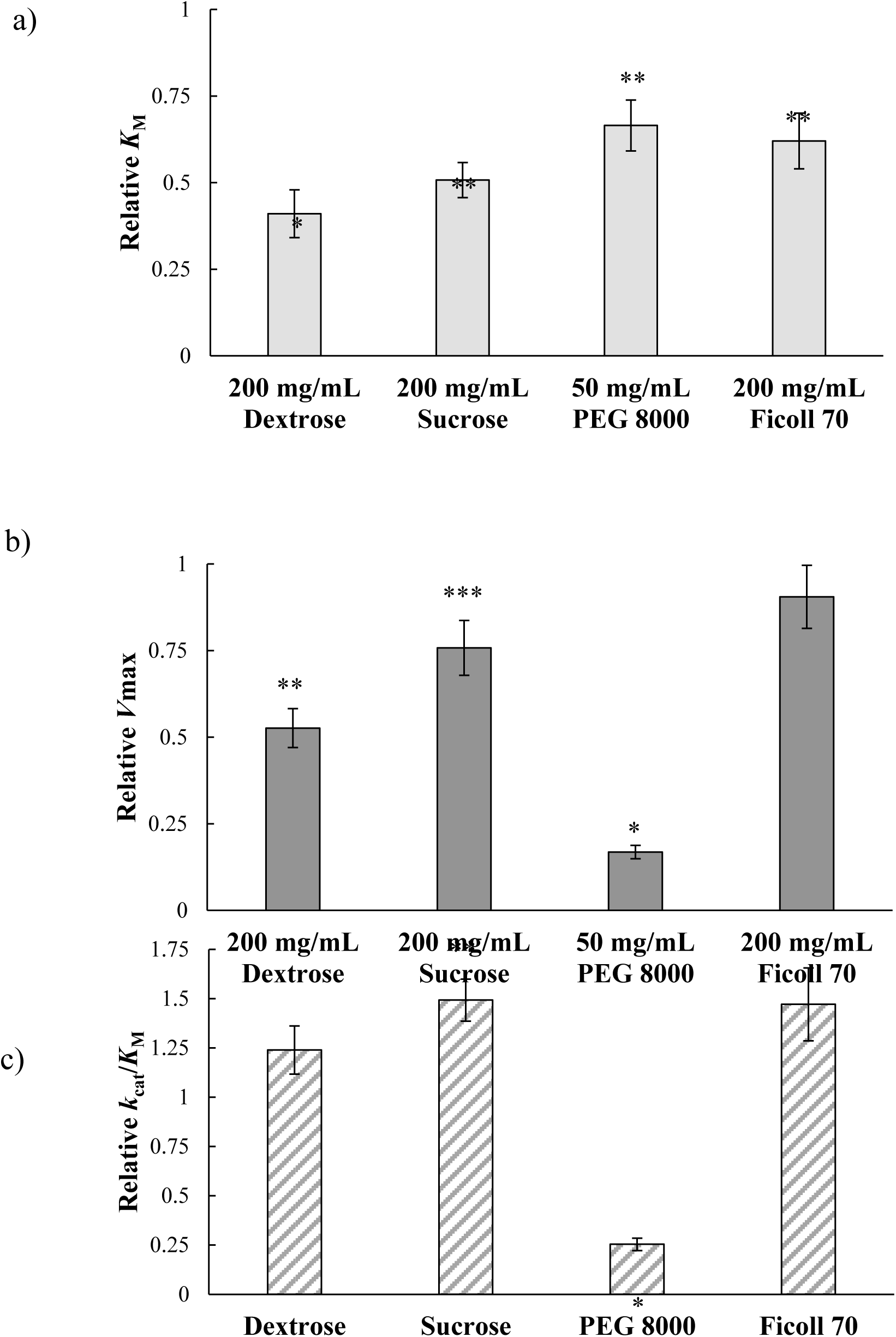
The kinetic parameters for Pro-AMP formation in the presence of various crowding agents: a) neutral crowding agents affect the *K*_M_ of Ec ProRS; b) Ficoll 70 was the only crowding agent resulting in no significant change in relative *V*_max_; c) sucrose enhanced relative *k*_cat_/*K*_M_ while PEG decreased relative *k*_cat_/*K*_M_. These studies were performed with 10 nM WT Ec ProRS and 0.125-1.00 mM proline. Reactions were incubated at 37 °C. A concentration of 200 g/L for PEG 8000 resulted in too significant of a reduction in catalytic activity to obtain accurate and consistent kinetic parameters such as *K*_M_ and *V*_max_. Error bars represent difference between two trials. Values for *K*_M_ and *V*_max_ were determined using a Lineweaver-Burk plot of a proline concentration variation assay (Fig. S1). Relative *K*_M_ and *V*_max_ displayed on the y-axis were normalized to the *K*_M_ and *V*_max_ determined under dilute condition. A two-tailed Student’s *t* test was performed to assess the significance of the difference between the relative kinetic parameters in the presence of crowding agents compared to dilute condition. The significance of the results of the *t* test are represented as follows: *p < 0.005, **p < 0.01, ***p < 0.05.

The observed alteration in binding (*K*_M_) and reaction rate (*V*_max_) of the formation of a prolyl-adenylate can be explained by excluded volume effect, where the overall accessible space is decreased in the presence of crowders resulting increase in local concentration of substrate.(49) Alternately, the crowders are also known to impact the conformational equilibrium favoring the substrate-bound conformation (49). In both cases, the stronger enzyme (E) – substrate (S) interactions would decrease the *K*_M_ of the proline activation, which is consistent with what observed in the present study. Furthermore, the stronger interaction would stabilize the E-S complex lowering its potential energy, which in turn would increase the activation barrier. The above scenario is consistent with the observed reduction in *V*_max_. However, a significant impact on *K*_M_ but not on *V_max_* in the presence of 200 mg/ml Ficoll 70 could be due to a subtle conformational change in the enzyme, which resulted in a slight decrease in *K*_M_. Similar observation was made earlier for *Enterobactin*-specific isochoramatic synthase (18).

### Impact of variable-sized polyethylene glycol on product formation

It was observed that the changes in Ec ProRS kinetic parameters (Fig. 2 and Table S2) did not correlate with the *R*_h_ (Å) of the crowding agents used in this study (Table 1). To examine the impact of the crowder size alone on proline activation (Pro-AMP formation), we eliminated the confounding effects of differences in chemical nature of crowders and performed end-point kinetics with variable-sized PEG molecules with final concentration of 200 g/L (Fig. 3; Supplementary Table S1). The main reason for choosing PEG is its simple chemical structure and the availability of variable-sized PEG with hydrodynamic radius comparable to the other crowders used in this study. A non-linear trend was observed between the Pro-AMP formation (nmol) and the hydrodynamic radius (*R*_h_) (Å) of variable-sized PEG (Fig. 3). It is apparent that when considering size alone, smaller-sized crowding agents have more impact on the catalytic function of Ec ProRS; the smaller the *R*_h_ of the given PEG, the less Pro-AMP was formed.

**Figure 3.**
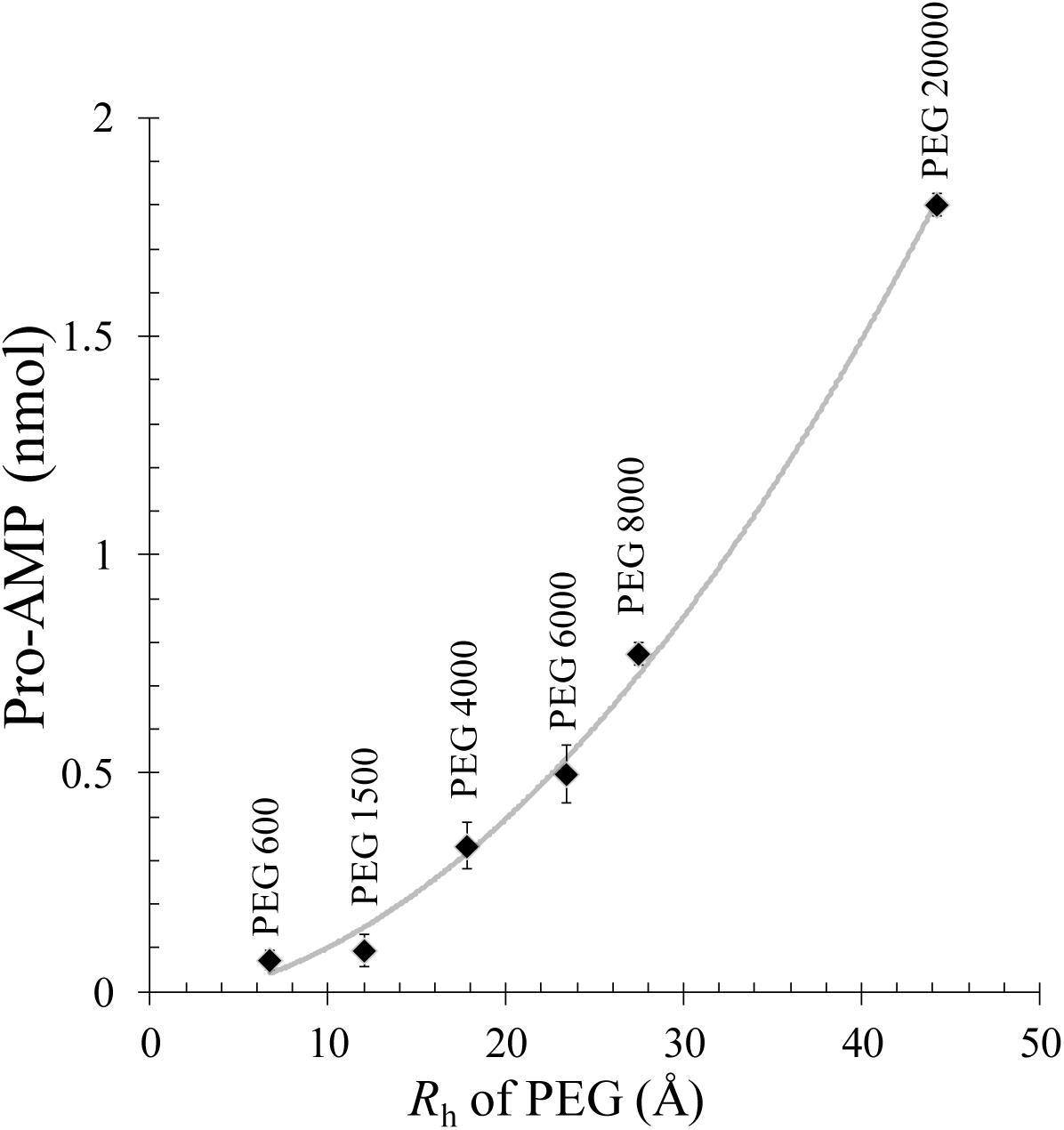
The hydrodynamic radii (Supplementary Table S1) of various length polyethylene glycol compared to formation of Pro-AMP (nmol). A concentration of 100 g/L PEG, 0.75 mM proline, and 10 nM WT Ec ProRS was used. Reactions were quenched at 20 minutes. PEG sizes from furthest left to right on graph are as follows: 600, 1500, 4000, 6000, 8000, 20000. Dilute condition had an average formation of 4.23 ± 0.04 nmol Pro-AMP.

One possible explanation of the observed trend in product formation is due to the poor accessibility of the enzyme active site in the presence of smaller PEG molecules. However, all PEG molecules used in this study are larger in size compared to the available space in the active site of Ec ProRS, which has an estimated volume of ∼500 Å^3^ (Fig. S2). Therefore, the reduction in product formation may not be due to the blockade of the enzyme active site. Rather, the observed higher impact caused by smaller PEG molecules hinted that these molecules may have “soft-interactions” with the enzyme. This is more likely since PEG is known to interact with proteins and capable of inducing conformational changes (50), which in the present scenario could also impact the conformational equilibrium. Taken together, observations from the size variation study suggest a lesser overall role of the excluded volume and steric hindrance (“hard-interactions”) in altering the enzyme function. On the other hand, these results indicate that the shift in conformational equilibrium could be one of the main mechanistic factors by which crowders may impact the function of Ec ProRS.

### Crowder-induced changes in intrinsic tryptophan fluorescence

To examine if “soft-interactions” are responsible for the shift in conformational equilibrium, intrinsic tryptophan fluorescence study was carried out in the presence of crowders. It has been well established that the intrinsic fluorescence of tryptophan is sensitive to the local environment and both fluorescence intensity and fluorescence wavelength are influenced by the extent of interactions of a tryptophan with its neighboring species (51). It is also known that if a tryptophan becomes more exposed to the surrounding hydrophilic solvent, there is a shift in the barycentric mean fluorescence wavelength (*λ*_bcm_) towards a higher wavelength (red-shift/Stokes shift) (51, 52). Conversely, if a tryptophan becomes less exposed to the polar solvent, the *λ*_bcm_ undergoes a spectral shift towards a lower wavelength (blue-shift/anti-Stokes shift) (51). Hence, changes in the tryptophan fluorescence emission properties serve as an excellent tool to assess “soft-interactions”, conformational changes, as well as substrate binding or denaturation (51). Therefore, the tryptophan fluorescence emission was monitored in the presence of crowding agents of different concentrations, sizes, and chemical natures. Ec ProRS is a dimer and has five tryptophan residues in each subunit (*vide infra*). Any observed effect of crowding on the intrinsic tryptophan fluorescence is supposed to be the combined effect resulting from the varied degrees of interactions of the five tryptophan residues with solvent/crowders molecules.

### Alteration in fluorescence emission intensity in presence of crowders

The intensity of fluorescence emission decreased as the concentration of crowders increased (Table 1 and Supplementary Fig. S3a-S6a). Here, dextrose was the exception, where 100 g/L had the greatest decrease and 300 g/L had the least effect on the decrease in emission intensity (Supplementary Fig. S3a). The greatest decrease in fluorescence emission intensity was produced by Ficoll 70 (76.7 %), followed by sucrose (51.9 %), then PEG 8000 (43.3 %); dextrose had the least effect on fluorescence emission intensity (6.52 % reduction) (Table 1). Although no definitive trend was observed between the size of the crowding agent and the percent reduction in fluorescence emission intensity, the larger crowders appeared to have more impact.

It is known that the quantum yield of tryptophan fluorescence decreases in a hydrophilic environment (53). The fluorescence quenching could be either due to collisions (dynamic quenching) or due to “soft-interactions” (static quenching) with the crowders and solvent molecules (53). In either case, the hydrophobic tryptophan fluorophore must have exposed to the surface, indicating moderate to strong interactions between Ec ProRS and crowding agents. This is supported by the fact that similar extent of reduction in fluorescence intensity was also observed for free tryptophan in the presence of molecular crowders. For free tryptophan, 300 g/L sucrose and dextrose resulted in 30% and 2.5% decrease in fluorescence intensity, respectively.

A mixture of crowding agents also had impact on fluorescence emission intensity. A combination of 100 g/L Ficoll 70 and 200 g/L sucrose resulted in the greatest decrease in emission intensity (Supplementary Table S3, Supplementary Fig. S7a-S9a). Solutions with a combination of sucrose and dextrose had lower percent of decrease in the emission intensity compared to other mixtures of two crowding agents (Supplementary Table S3). There was also a correlation between the percentage of emission intensity reduction and the total concentration of crowding agents, with higher total concentrations of crowders resulting in a higher percentage reduction than lower concentrations (Supplementary Table S3). A closer analysis revealed that the impact of the mixture of two crowding agents on the fluorescence intensity (Table 1 and Supplementary Table S3) is additive suggesting that crowding agents are acting in a non-competitive manner.

### Shift in barycentric mean fluorescence wavelength in presence of crowders

All crowding agents, except for dextrose, showed increasing *λ*_bcm_ (red-shift) with increasing concentrations of crowding agents; Ficoll 70 resulted in the largest increase in *λ*_bcm_ (Table 1 and Supplementary Fig. S3b-S6b). Dextrose, being the exception, resulted in a Δ*λ*_bcm_ that was slightly negative compared to dilute conditions (Table 1). There was no distinct trend between the *R*_h_ of these four crowding agents and Δ*λ*_bcm_ (Table 1). However, it took approximately twice the concentration of sucrose to achieve the same increase in *λ*_bcm_ as observed for Ficoll 70. Since the resultant fluorescence emission spectra is the sum total of emissions from all the ten tryptophan residues in the dimeric Ec ProRS, we can only say that the crowding effect led to a net increase/decrease in tryptophan exposure to the solvent (51). The observed changes in *λ*_bcm_ indicate considerable interactions between tryptophan residues and crowder molecules, capable of inducing a change in the conformation of Ec ProRS. It is important to note that for free tryptophan, Δ*λ*_bcm_ was ≤1 nm in 300 g/L sucrose and it remained practically unchanged in the presence of 300 g/L dextrose.

Taken together, the alteration in *λ*_bcm_ and the fluorescence intensity in presence of crowding agents is an indicative of “soft-interactions” between these crowders with Ec ProRS, which could induce a change in the conformational equilibrium further impacting its function. Similar observation has been made for other protein systems, where protein folding, and conformational equilibrium were affected by molecular crowding (20, 21, 54–56). To obtain a molecular-level understanding of how a crowding agent alters an enzyme’s conformational equilibrium, we took a closer look at the impact of the two crowding agents, dextrose and sucrose, on the Ec ProRS conformation and dynamics using molecular dynamics simulations.

### Molecular dynamics simulations to probe the crowder-induced conformational dynamics

**S**imulations of the dimeric Ec ProRS were performed by maintaining the same conditions as used in fluorescence experiments. Enzyme systems were built using the following three conditions: i) in water or dilute, ii) in the presence of 200 g/L dextrose, and iii) in the presence of 200 g/L sucrose. As a representative system, the sucrose containing solvated system is shown in Fig. 4. In the present study, only dextrose and sucrose were chosen because i) they have considerable impact on kinetic parameters (Fig. 2), and ii) due to the simplicity in building the solvent box with these two crowding agents. To ascertain any changes in collective domain motions, the essential dynamics of the protein were studied by principal component analysis (PCA). In addition, solvent accessible surface area (SASA) calculations were carried out to explore the changes in the conformation of the protein over the course of entire simulations.

**Figure 4.**
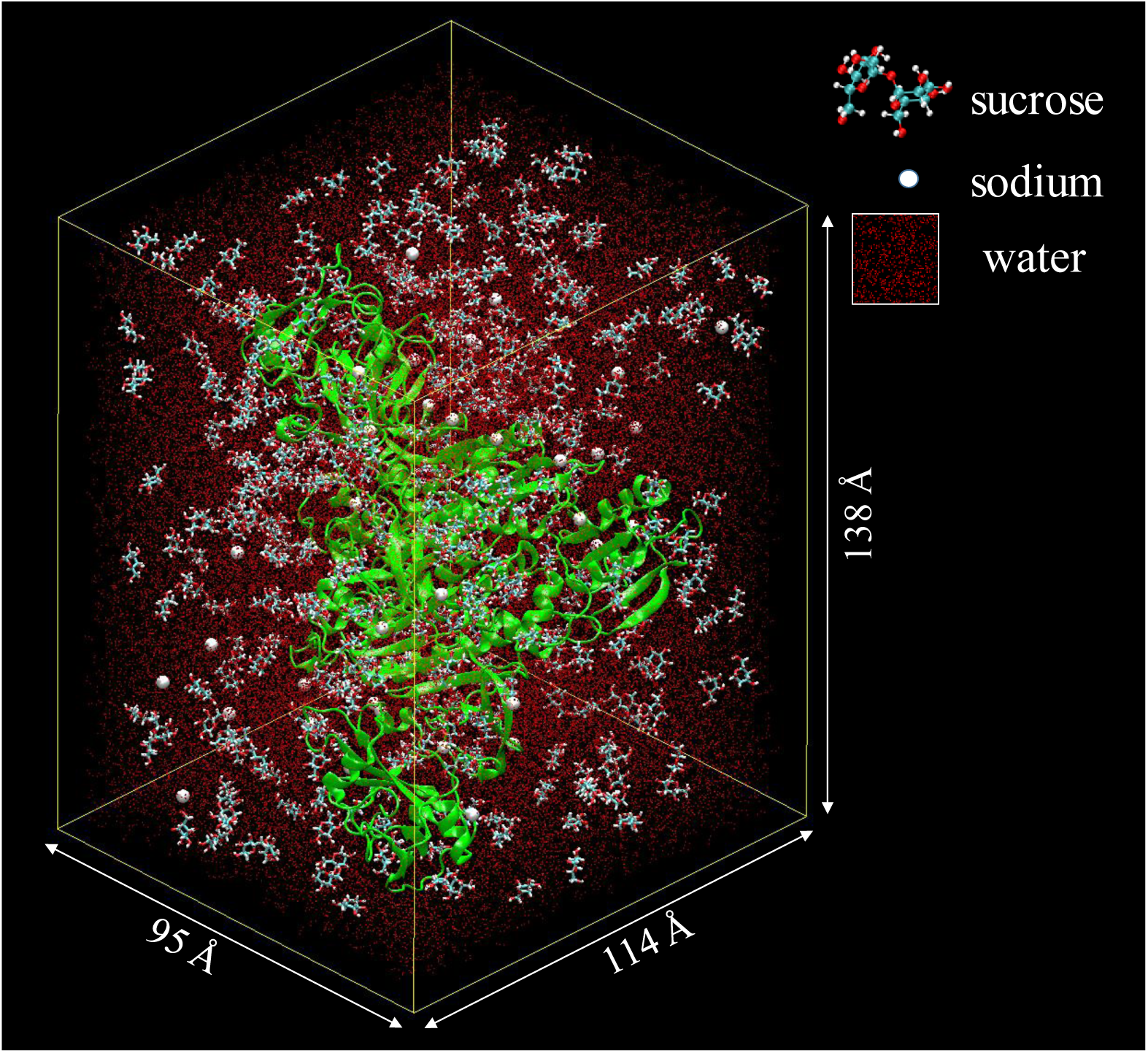
The dimeric Ec ProRS in a solvent box containing 200 g/L sucrose molecules used for 70 ns MD simulations. The setup resulted in a total number of 766 sucrose molecules, 31100 water molecules, and 42 sodium ions. The total number of atoms in the system were 128313.

The SWISS-MODEL homology server generated two models (one per monomer unit), each of which had Global Model Quality Estimation (GMQE) scores of 0.78 and coverage of > 95% (data not shown). The resulting dimeric structure was analyzed using Procheck (57, 58). A total of 87.0% of the residues were in the most favored regions of the Ramachandran plot, 11.6% in additional allowed regions, and only 5 of (0.5%) residues in disallowed regions (Supplemental Fig. S10). These results ensured that the model structure of the dimeric Ec ProRS is of reliable quality and suitable for MD simulations.

### Impact of crowders on protein conformational distribution

Each subunit of Ec ProRS has three distinct domains – editing domain (also known as insertion or INS domain), the central catalytic domain (CD), and the anticodon binding domain (ABD) (Fig. 5a). The two crowding agents, dextrose and sucrose, were observed to have significant impact on the motion of INS domain (Fig. 5). In the optimized structures, the protein segment with residues 313 to 322 of INS domain was found to be located about 6 Å away from the small helical segment (residues 84 to 93) of the central CD (Fig. 5a). The proximity between the above two segments was observed for all 3 systems indicating that the “closed” conformation occurs at the start of each simulated system. During 70 ns simulation, INS domain was observed to move away from CD forming an “open” state in the absence of any crowding agent (Fig. 5a). The extent of opening was measured from the displacement of the INS domain relative to the CD. The inter-domain distance was defined by the C_α_-C_α_ distance of residues Q88 of CD and P318 of INS. From the evolution of the structure during 70 ns MD simulations, it is evident that the two domains, INS and CD, moved apart by ∼ 25 Å in the absence of any crowding agents (Fig. 5b). On the other hand, smaller domain displacement (< 15 Å) was observed in the presence of crowding agents indicating that crowders appeared to have favored the “closed” conformation. The distribution of conformations in the three systems was studied and discussed below.

**Figure 5.**
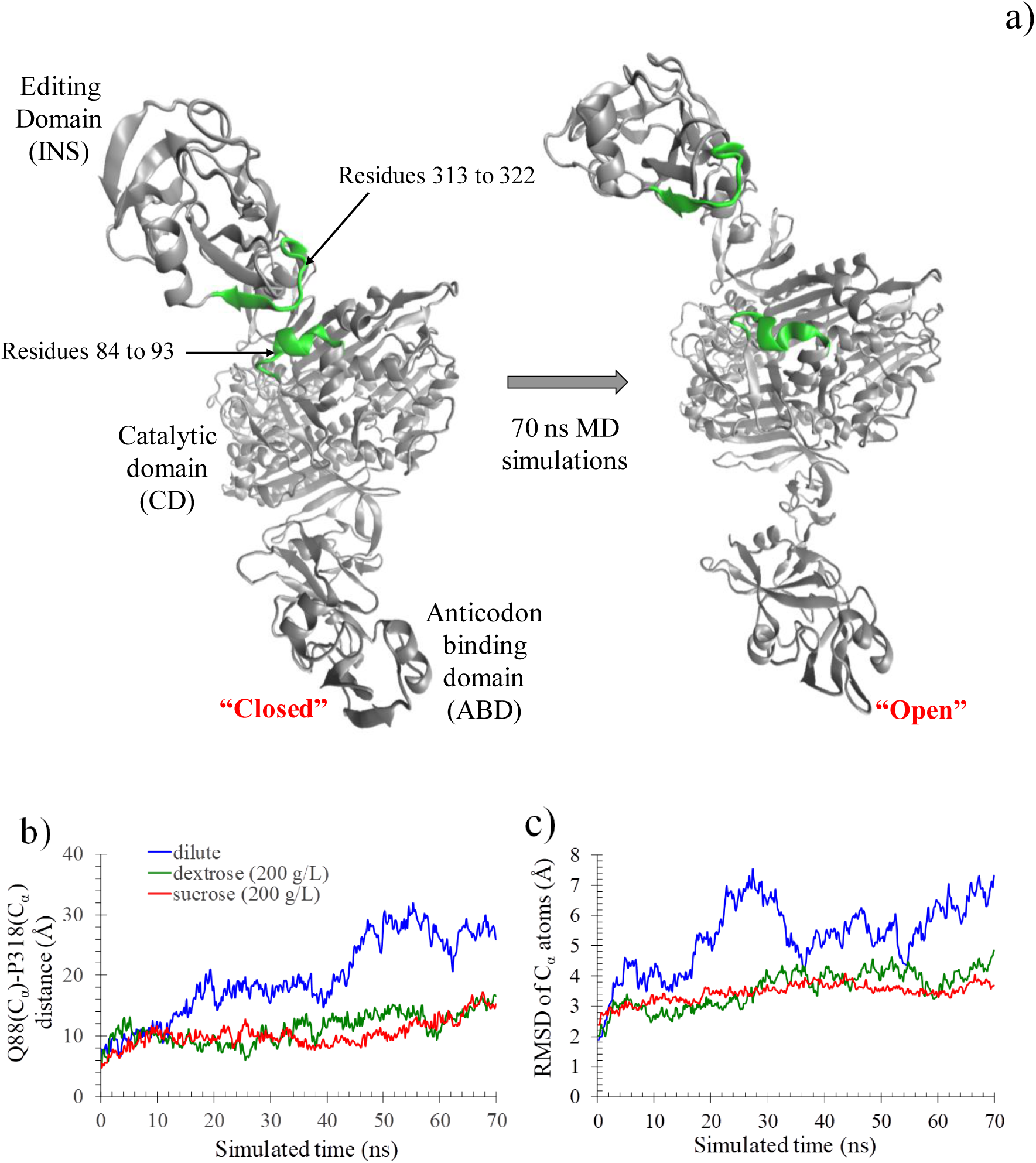
a) Structures of Ec ProRS in “closed” and “open” states. The editing domain residues (residues 313 to 322) and catalytic domains residues (residues 84 to 93) are shown in green color; b) evolution of the distance (in Å) between C_α_ atoms of Q88 and P318 over the simulation period of 70 ns in three systems: dilute condition, dextrose (200 mg/ml), and sucrose (200 mg/ml); c) the backbone root-mean-square-deviation (RMSD) of each frame calculated from their starting conformation over the simulation period of 70 ns for the three systems.

The ratio of “open” to “closed” conformations was measured from the MD trajectories (Table 2, Fig. 5). The conformations were defined as “closed”, where the inter-domain distance (i.e. Q88(C_α_)-P318(C_α_)) was ≤ 10 Å. On the other hand, if the inter-domain distance was ≥ 15 Å, the conformations were considered as “open”. The “intermediate” conformations had the Q88(C_α_)-P318(C_α_) distance between 10 – 15 Å. Analysis of the simulated trajectories revealed that the “closed” conformational state is predominant (49 %) in sucrose containing system, while a reverse trend was observed in the absence of any crowder molecule with ∼76 % conformations observed in the “open” state. The dextrose containing system appears to favor the “intermediate” state more compared to the dilute system (Table 2). Also, the plot of Q88(C_α_)-P318(C_α_) versus simulated time demonstrated that the compact conformation was preferred in sucrose and dextrose and the extended conformation in dilute condition (Fig. 5b).

**Table 2.**
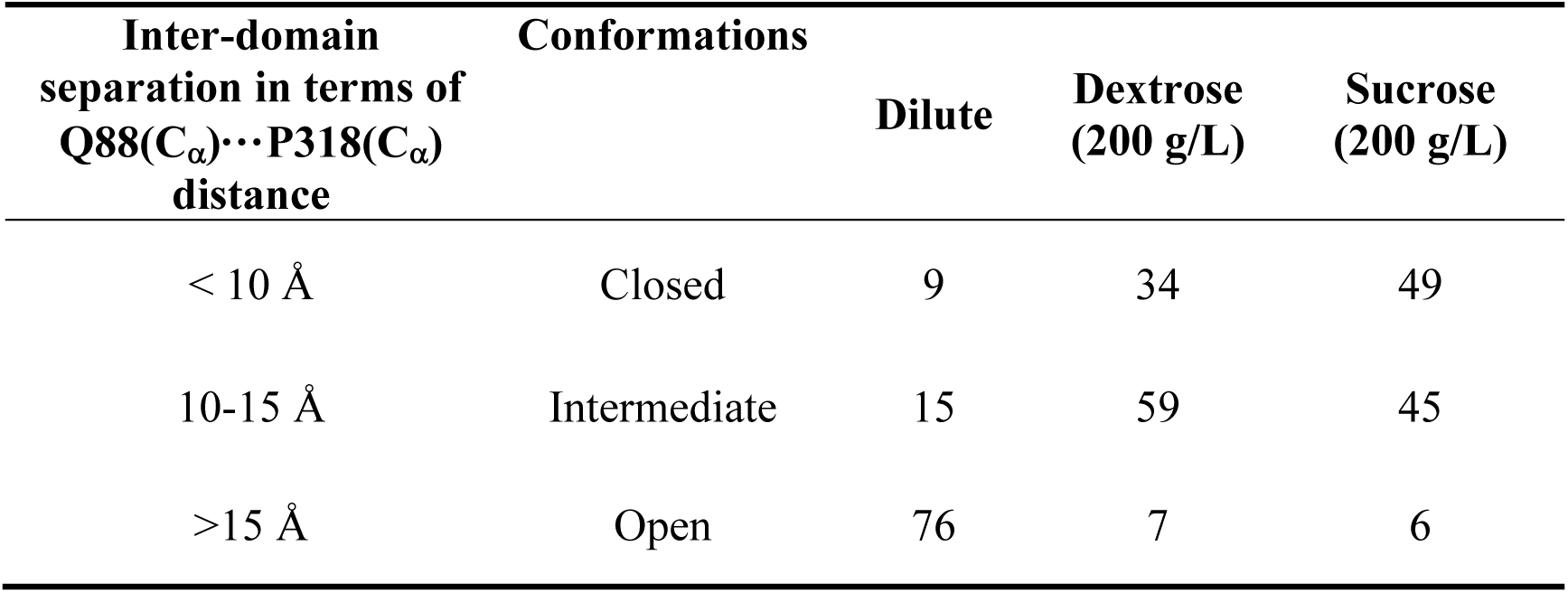
Change in conformational ensemble along the 70 ns trajectory of MD simulations

It is to be noted that the same concentration of sucrose and dextrose was used in MD simulations and kinetic assays. Thus, the changes in the simulated dynamics of the substrate-free enzyme are expected to be correlated to the kinetic assays and hence could provide insights into the observed catalytic changes. The shift in the conformational equilibrium from “open” to “closed” state in the presence of dextrose and sucrose correlates very well with the increased binding affinity for the substrate. As certain segments of editing domain were found to remain much closer to the catalytic domain in the presence of both crowders (Fig. 5a), the substrate-bound state would be favored, which is consistent with the decreased *K*_M_ as observed experimentally from the kinetic assays.

The plots of backbone RMSDs of each conformation (of the evolved molecules during the 70 ns simulations under different conditions) with respect to the initial equilibrated structure are shown in Fig. 5c. In each case, the RMSDs were computed from the optimized starting structure of the solvated dimeric enzyme. The overall effect of these crowders on backbone fluctuation displayed a strong damping due to crowders. The evolution of RMSDs demonstrated that the backbone RMSD fluctuated by only 3 – 4 Å in presence of crowders, as compared to the dilute solution where a 7 Å fluctuation was noted. As the backbone fluctuation is predominantly due to the motion of INS domain, the damping in backbone fluctuation correlates well with the “open” to “closed” population shift in the presence of dextrose and sucrose. The crowders-induced damped fluctuation of the backbone further indicates the presence of “soft-interaction” between protein side chains and crowders.

### Changes in solvent accessible surface area (SASA) and radial distribution function (RDF) in crowded environments

If there is a shift in conformational distribution due to the presence of “soft-interactions” between crowders and protein side chains, it is expected that the SASA of the full protein and individual residues would change. Therefore, the MD simulation data was analyzed to explore the sucrosx1e- and dextrose-induced solvent accessibility of individual tryptophan residues. As mentioned earlier, there are five tryptophan residues in each subunit of the dimeric Ec ProRS, the locations of these residues are displayed in Fig. 6a. The SASAs of each tryptophan residue were calculated for every 10 ns (of the total 70 ns MD simulation) in three different conditions - dilute, dextrose and sucrose (Fig. 6b). The alteration in tryptophan residues’ SASA was noticed in the presence of sucrose and dextrose. In particular, the SASA values of W375 of INS domain exhibited a spectacular change – the SASA changing from ∼25 Å^2^ in the dilute to ∼70 - 90 Å^2^ for the dextrose and sucrose, respectively. In contrast, a decrease in SASA was observed for W86 in the presence of crowding agents (Fig. 6b). The alteration in SASA values confirmed the crowder-induced conformational change in the enzyme.

**Figure 6.**
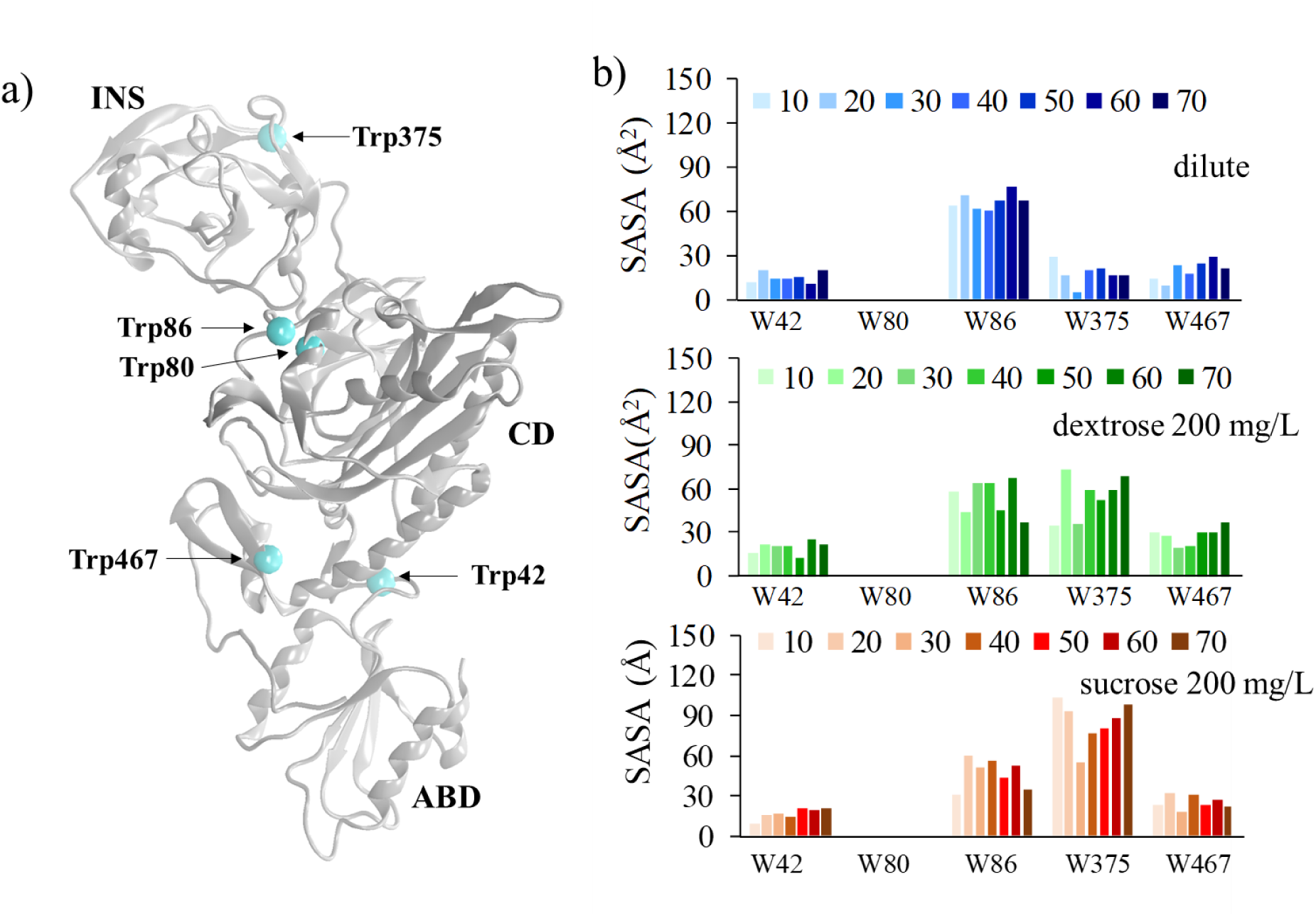
The effect of crowding agents on “soft-interactions”: a) location of tryptophan residues shown as cyan beads on wild-type Ec ProRS monomer. The monomer is shown in “NewCartoon” and the three distinct domains of Ec ProRS are labelled as INS (editing domain), CD (catalytic domain), and ABD (anticodon-binding domain); b) computed SASA for tryptophan in the dilute condition (blue), 200 g/L dextrose (green), and 200 g/L sucrose (orange) are shown as histograms. All SASA were tracked every 10 ns of the simulation.

To further examine the effect of “soft-interactions” between the crowder molecules and protein side chains, radial distribution function (RDF) of water, dextrose, and sucrose with respect to the two tryptophan residues, namely, W86 and W375 were computed. These residues were observed to have higher SASA values (Fig. 6b). The RDF plot shows the probability of finding water or crowder molecules within the spherical region surrounding the tryptophan residue (Fig. 7a-b). The center of the sphere represents the geometric average of all atoms of the respective tryptophan residue. The RDF plot shows a flat curve for water, but distinct peaks for both dextrose and sucrose, which indicates higher probability of finding crowder molecules within a certain distance of the tryptophan, relative to water. It is to be noted that the water molecules in these simulations greatly outnumbered the crowder molecules – for each dextrose molecule there were 31 water molecules, while sucrose-to-water molecule ratio was ∼ 1: 40. This observation coupled with the increased exposure of the tryptophan to the solvent (i.e. SASA) confirmed the presence of “soft-interactions” between protein and crowder molecules (dextrose and sucrose).

**Figure 7.**
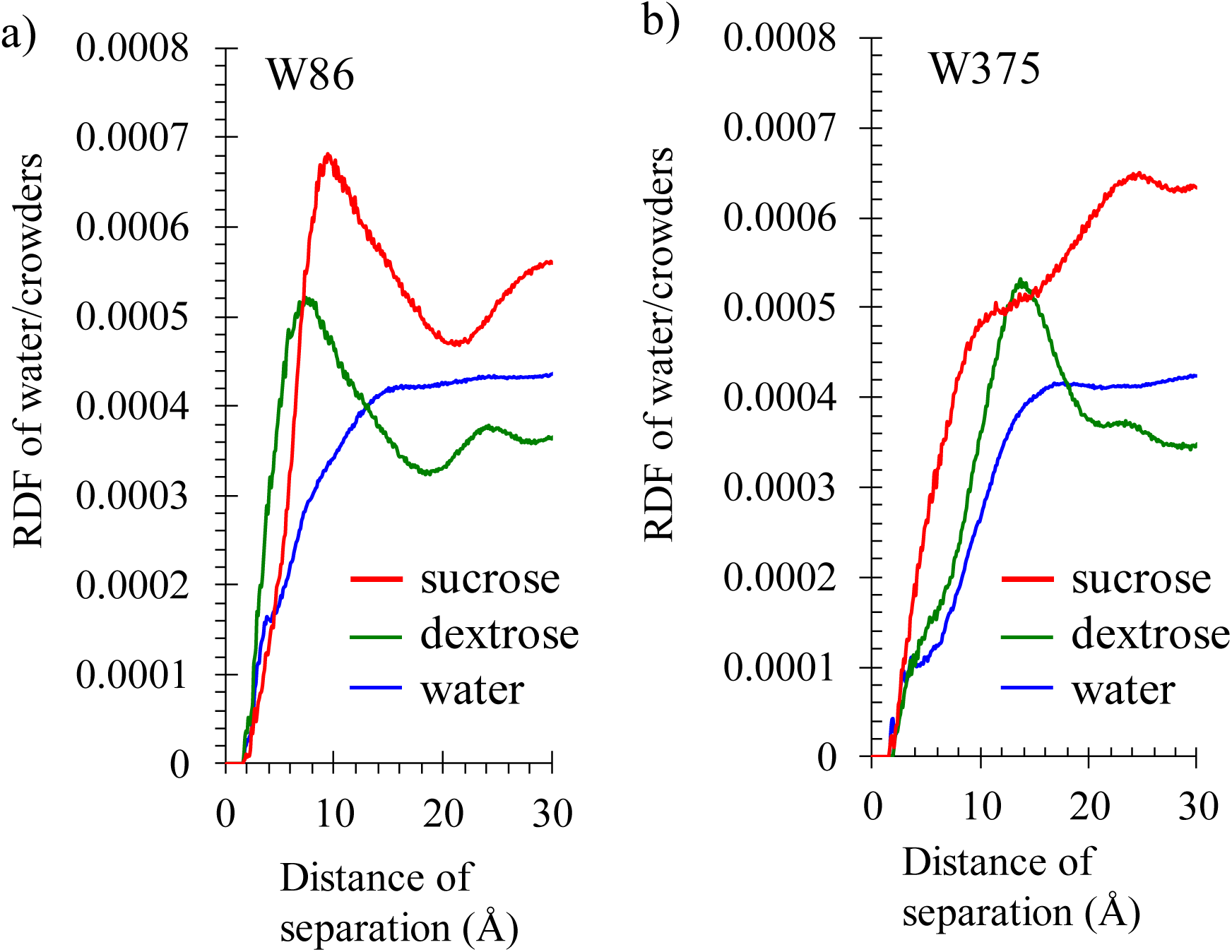
Radial distribution function of water, dextrose, and sucrose molecules with respect to a) Trp86 and b) Trp375 of Ec ProRS, computed using the last 60 ns MD simulation data. The x-axis represents the distance of separation between the center of mass of the tryptophan residue and the center of mass of water, dextrose, and sucrose in dilute, 200 g/L dextrose, and 200g/L sucrose, respectively.

### Potential energy surface calculation to probe soft-interactions between tryptophan and sugar molecules

The higher RDF and SASA values of W375 and W86 indicate a stronger interaction between the tryptophan and dextrose/sucrose than water. The dispersion interaction or CH/π interaction between carbohydrates with aromatic amino acid residues have been reported earlier (59, 60). To examine if dispersion interaction is prevailing, the potential energy surfaces (PES) of the three systems were computed using approximate quantum chemical formalism (SCC DFTB-D module of CHARMM program) (29, 30, 47). The observed PESs depict the strength of the interaction of the indole ring of the tryptophan with water, dextrose, and sucrose in terms of the depth of the potential energy well. The calculated potential energies are plotted against the distance between the centroid of the benzene ring of indole and the centroid of the pyranose (sugar) molecule. In the case of water, the oxygen atom is used in the place of pyranose. The comparison of potential energy wells of the three systems is shown in Fig. 8. The lowest point of the potential energy well was found to be only −0.2 kcal/mol occurring at 3.6 Å for water. In contrast, much stronger interactions were noted for both dextrose and sucrose systems (−6.2 kcal/mol and −6.9 kcal/mol occurring at 3.1 Å, respectively; Fig. 8). These results indicate the presence of ∼ 30 − 35 times stronger interactions between the sugar ring of dextrose/sucrose and the indole ring of the tryptophan compared to the interaction between water and tryptophan. Therefore, the PES calculations revealed a molecular basis of how “soft-interactions” are playing a role in altering the conformational equilibrium of Ec ProRS.

**Figure 8.**
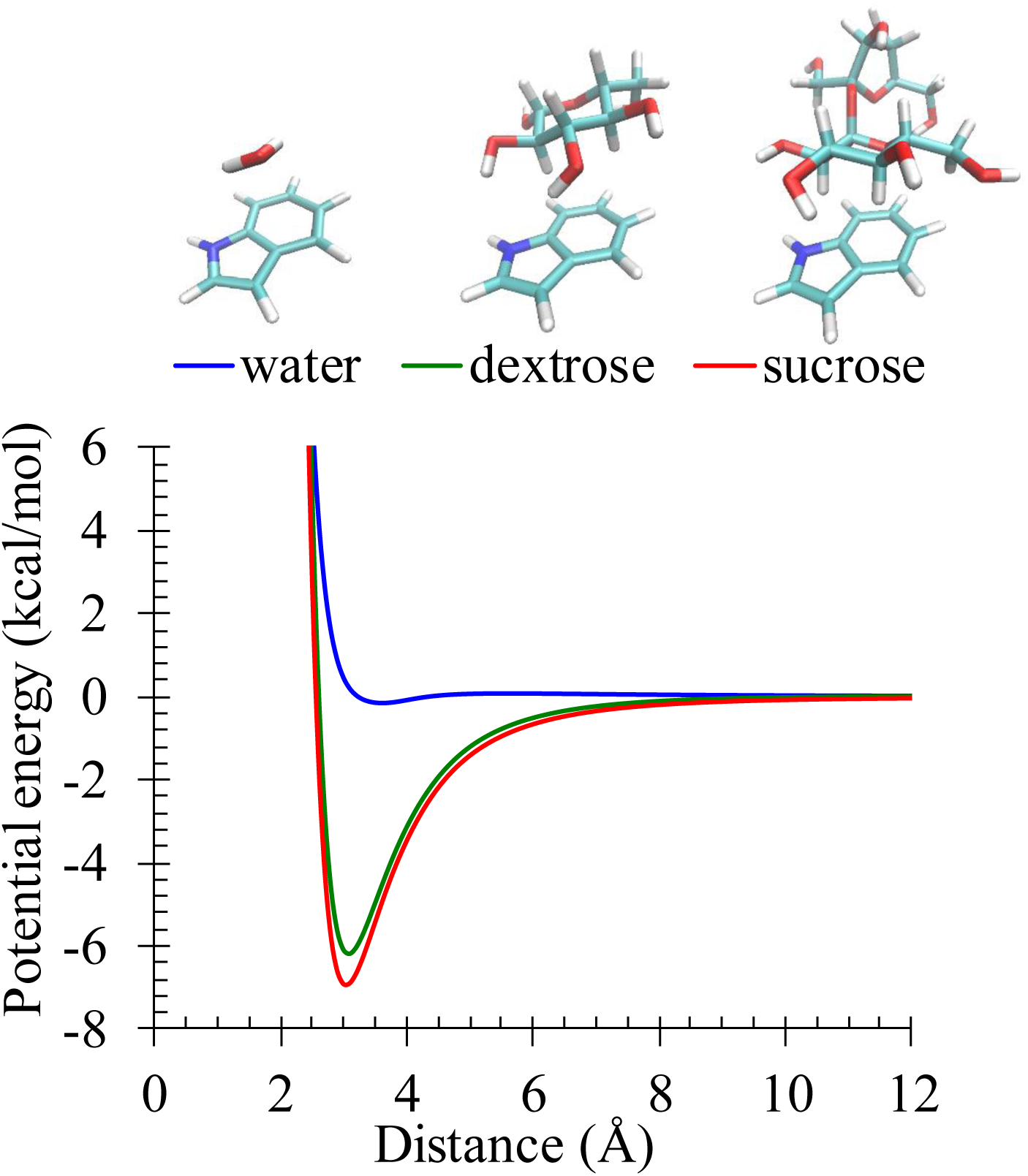
The gas-phase potential energy of interaction between the tryptophan with water, dextrose, and sucrose, computed using approximate quantum chemical formalism (SCCDFTB-D module of CHARMM program). The intermolecular distance is the distance between the centroid of the benzene ring of indole and the centroid of the pyranose (sugar) molecule. In the case of water, the oxygen atom is used in the place of pyranose.

### Crowder-induced alterations in coupled-domain dynamics

To explore the impact of crowding agents on the collective movements of the protein backbone, the essential dynamics of Ec ProRS in dilute and crowded conditions were examined by computing their principal components of motions. In particular, the dynamical changes in the first three principal components were examined for each system. The principal component analysis (PCA) revealed an alteration in the collective domain dynamics in the presence of dextrose and sucrose (Fig. 9). The editing and catalytic domain were observed to move in an anticorrelated manner. However, their motions were restricted in the presence of dextrose and sucrose (Fig. 9, left to right). In particular, the collective dynamics of the editing domain and the key secondary elements at the interface of the editing domain and the catalytic domain, which includes proline-binding loop (residues 195 to 210), the helix-loop-beta (residues 305 to 322) and the helix (residues 84 to 93) were altered in the crowded environment (Fig. 10). An increased extent of compactness surrounding these structural elements, which comprise the active site pocket of Ec ProRS, was observed in the presence of dextrose and sucrose. Earlier we have reported that the coupled-dynamics between distant structural elements are important for efficient catalysis by Ec ProRS.(6, 7) In the present study, a significant change in the dynamics of PBL was noted. As observed in the dilute condition, the movement of PBL (red to blue ribbon in Fig. 9 and 10) was more pronounced compared to that in the presence of dextrose and sucrose. Other structural elements also moved to a lesser extent due to crowding creating a thermodynamic shift in the conformational equilibrium and thus impacting the enzyme function. The observed decrease in *V*_max_ can also be rationalized as the limited opening of the active site pocket would lead to reduced frequency of effective collisions with substrates as well as decreased product release. Moreover, when considering the dimeric Ec ProRS, the separation between INS domains of the two subunits were also found to be altered in the presence of crowders (Fig. 11). The distance between the two INS domains decreased considerably in the presence of dextrose and sucrose, which could also alter the substrate binding in the presence of crowders. Therefore, results of MD simulations support the observed changes in kinetic parameters in the presence of crowders.

**Figure 9.**
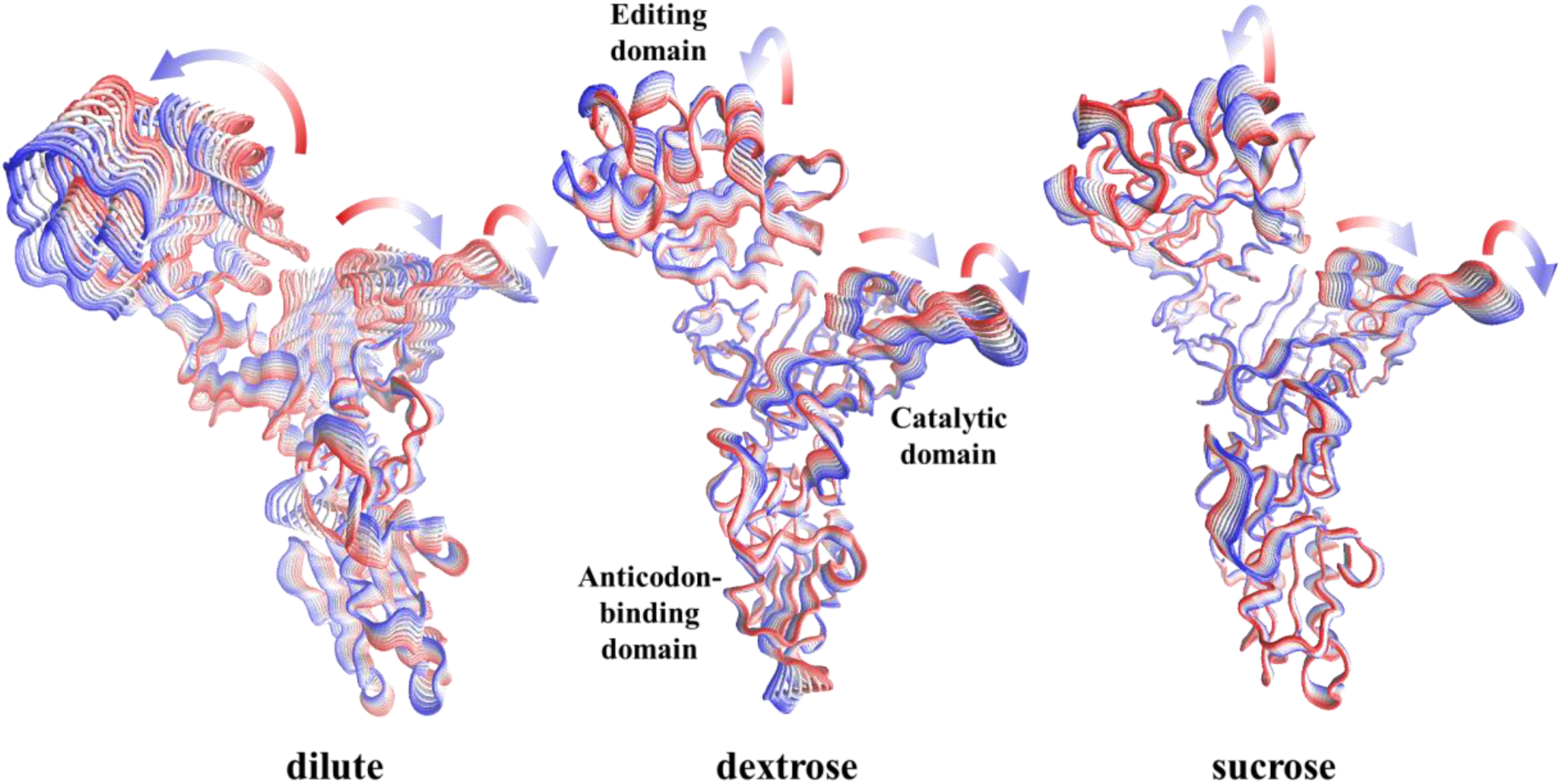
Principal component analysis of the dilute, dextrose (200 g/L), and sucrose (200 g/L) systems using the 70 ns simulation data. The red-colored ribbons depict the position of the backbone C_α_ atoms of Ec ProRS at the beginning and the blue-colored ones represent the same at the end of the 70 ns simulation.

**Figure 10.**
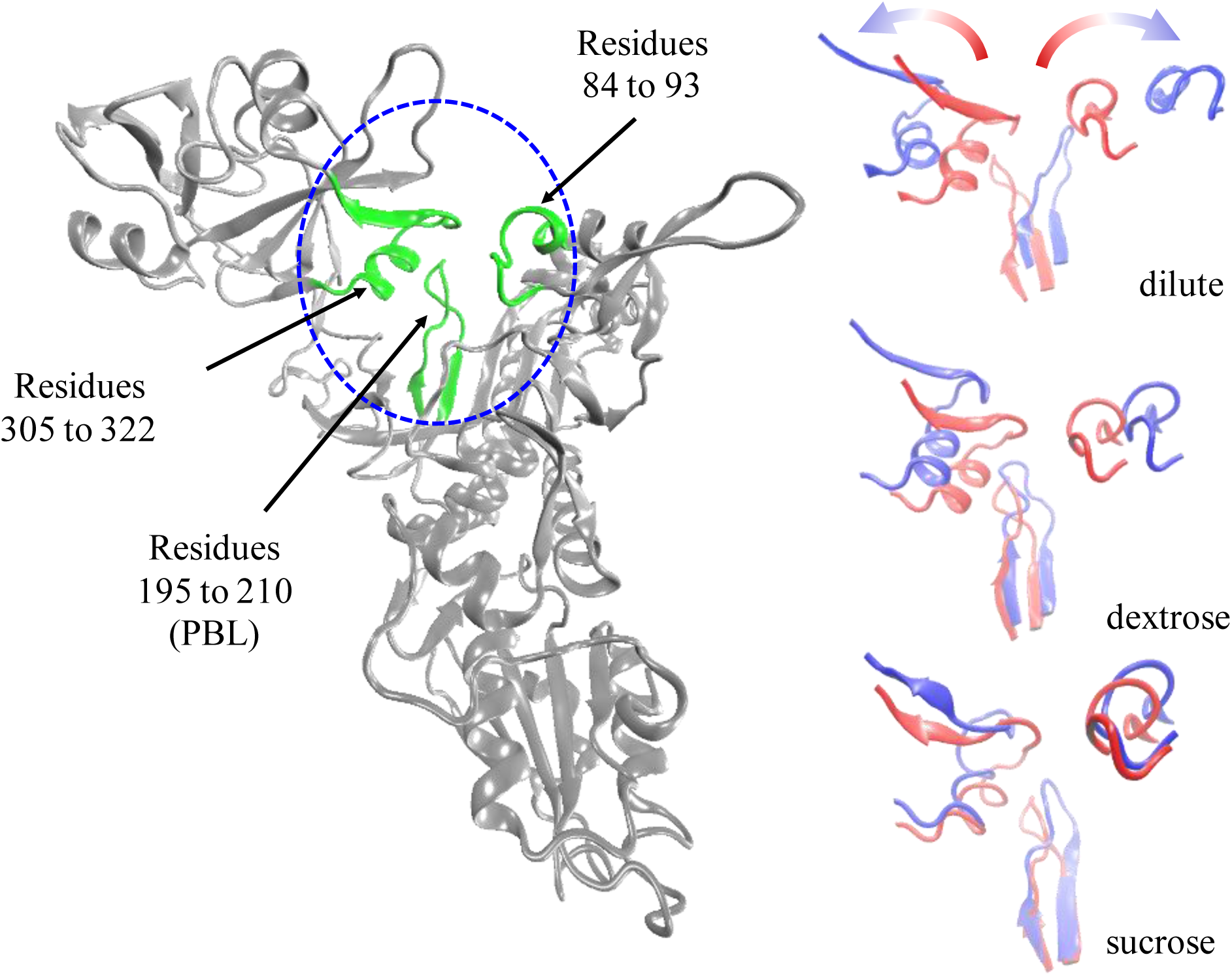
The conformational change observed during 70 ns MD simulations of Ec ProRS in dilute, 200 g/L dextrose, and 200g/L sucrose. The monomeric structure with the secondary structural elements surrounding the catalytic pocket, highlighted in green, is shown on the left. The displacements of these secondary elements during 70 ns simulations are shown on the right, where red and blue colors representing the starting and the ending conformations, respectively.

**Figure 11.**
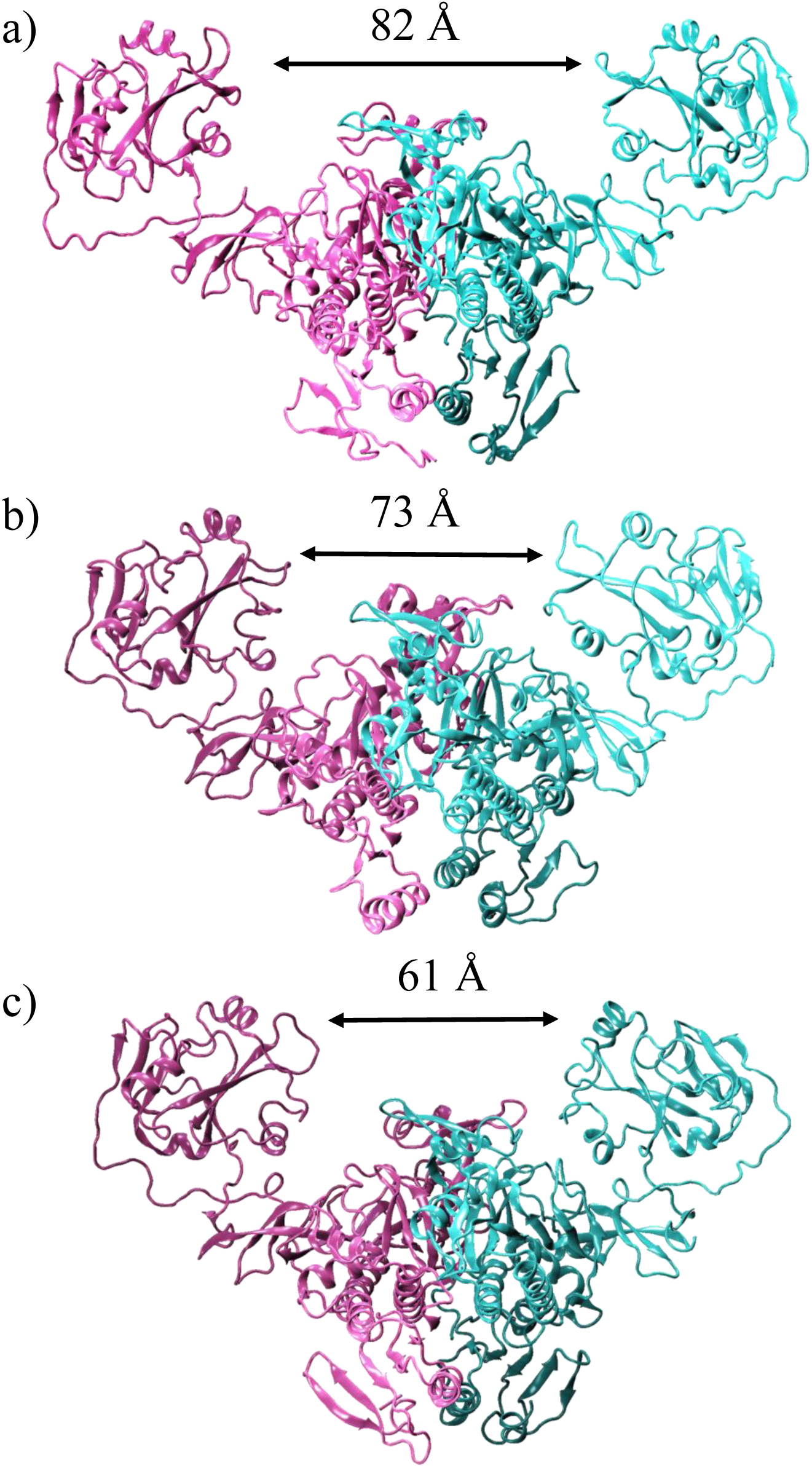
Varying degrees of compactness in the dimeric structure of Ec ProRS in the crowded environments: a) dilute; b) 200 g/L dextrose; and c) 200g/L sucrose.

## CONCLUSIONS

The present study showed that molecular crowding impacts the conformational equilibrium and dynamics of Ec ProRS, which leads to an alteration in the catalytic function. The simulations results demonstrated that stable interactions between crowders and tryptophans on the protein surface do exist, which is consistent with the reduction in fluorescence intensity and shift in the *λ*_bcm_. Approximate quantum chemical computations demonstrated that the interaction between the indole ring of the tryptophan and dextrose/sucrose is sufficiently stronger, attractive, and medium-range (3 – 8 Å) compared to the interaction with water. Furthermore, MD simulations and PCA confirmed that these “soft-interactions” alters the dynamics and hence changes the thermodynamic equilibrium between “closed” and “open” states. The observed favorability of the “closed” conformation in the presence of dextrose and sucrose is also consistent with the tighter proline binding and reduced *V_max_*. Although, only the interaction between the side chain of tryptophan and crowder molecules are probed in the present study, it must be noted that side chains of other amino acids especially, those on the surface must also have “soft-interactions” with the crowders molecules. These “soft-interactions” would result in altered conformational distribution and dynamics, which would ultimately have significant impact on catalytic function of Ec ProRS.

While information regarding the impact of macromolecular crowding is emerging every day, there remains many questions about its mechanism of action and variable effects it has on structure and function of modular proteins. The present study has found that the concentration, size, and the chemical nature of crowders have noticeable impact on Ec ProRS function. These variables are known to influence enzyme function through steric hinderance (“hard-interactions”) and/or “soft-interactions”. The present study has added additional insights into the molecular mechanism of crowding effects on the function of modular enzymes like ProRSs. It has demonstrated that crowding agents alter the conformational and dynamical properties of multi-domain Ec ProRS through “soft-interactions”. This resulted in a change in conformational distribution; the “closed” compact conformation was favored over the “open” extended conformation. In summary, this study is expected to serve as a valuable guide for predicting crowding-induced changes in conformation and dynamics of modular enzymes in cellular environment. This present study also demonstrated the significance of considering the macromolecular crowding effect while probing mechanism of catalytic activity of modular enzymes.

To our knowledge, this is the first report of crowder-induced shift in conformational ensemble in modular ProRS, an important member of AARS family. AARSs have emerged as important targets for anti-infective drug development because of its essential role in protein biosynthesis.(61–64) The documented effects of macromolecular crowding on conformation and function may have future implications in drug design. The present study suggested that the screening of potent drug molecules for pathogenic enzymes requires a thorough investigation of the impact of macromolecular crowding on the catalytic function of these enzymes. Finally, this study has exemplified the need for a multifaceted investigatory approach involving kinetic, spectroscopic, and computational techniques to elucidate the overall effect of macromolecular crowding on enzyme structure and function. Further studies with crowding agents mimicking the intracellular biomolecules and metabolites are needed to be carried out employing both experimental and computational approaches as it provides a greater insight into the structure-dynamic-function relationship of multi-domain enzymes in the intracellular environment.

## ACKNOWLEDGEMENTS

We would like to acknowledge UW-Eau Claire Material Sciences Department and the Blugold Supercomputing Cluster (BGSC) for the use of their facilities and equipment. We also acknowledge the help of the Iowa State University College of Liberal Arts and Sciences for providing access to the high-performance computing facilities, as well as the support of Dr. Walter Moss. This work was supported in part by National Institute of Health [grant number 1R15GM117510-01 (S.H. and S.B.)], XSEDE [grant number MCB110173 (S.H.)], and by the Office of Research and Sponsored Programs of the University of Wisconsin-Eau Claire, Eau Claire, WI.

## NOTES

The authors declare no conflict of interest.

## ABBREVIATIONS USED

AARSs: aminoacyl-tRNA synthetases
ProRS: prolyl-tRNA synthetases
WT: wild-type
Ec: *Escherichia coli*
PEG: polyethylene glycol
MD: molecular dynamics
*λ*_bcm_: barycentric mean fluorescence wavelength
VMD: visualize molecular dynamics
PCA: principal component analysis
SASA: solvent accessible surface area
DCCM: dynamical cross-correlation matrix.

## SUPLEMENTAL INFORMATION

Hydrodynamic radii of PEGs used in size variation study (Table S1); Kinetic parameters (Table S2); Intrinsic tryptophan fluorescence in the presence of a combination of crowding agents (Table S3); Lineweaver-Burk plot (Figure S1); A cartoon representation of the monomeric Ec ProRS showing the dimension of the active site pocket (Figure S2); Intrinsic tryptophan fluorescence emission spectra of Ec ProRS in the presence of varying concentrations of crowding agents (Figures S3a-S9a); Plots of *λ*_bcm_ versus concentrations of crowding agents (Figures S3b-S9b); Ramachandran plot of the homology model structure of Ec ProRS dimer (Figure S10).

## AUTHOR CONTRIBUTIONS

L.A., H.S., and S.H. designed and performed the wet-lab experiments; L.A. and R.A. performed fluorescence studies; R.A., Q.H., and S.B. designed and performed the computational experiments; L.A., R.A., S.H. and S.B. analyzed the data and wrote the manuscript.

## REFERENCES

1. Delarue, M., and D. Moras. 1993. The aminoacyl-tRNA synthetase family: modules at work. Bioessays 15:675–687.

2. Ibba, M., and D. Soll. 2000. Aminoacyl-tRNA synthesis. Annu. Rev. Biochem. 69:617–650.

3. Cusack, S., A. Yaremchuk, and M. Tukalo. 2000. The 2 Å crystal structure of leucyl-tRNA synthetase and its complex with a leucyl-adenylate analogue. EMBO J. 19:2351–2361.

4. Tukalo, M., A. Yaremchuk, R. Fukunaga, S. Yokoyama, and S. Cusack. 2005. The crystal structure of leucyl-tRNA synthetase complexed with tRNALeu in the post-transfer-editing conformation. Nat. Struct. Mol. Biol. 12:923–930.

5. Crepin, T., A. Yaremchuk, M. Tukalo, and S. Cusack. 2006. Structures of two bacterial prolyl-tRNA synthetases with and without a cis-editing domain. Structure 14:1511–1525.

6. Sanford, B., B. Cao, J. M. Johnson, K. Zimmerman, A. M. Strom, R. M. Mueller, S. Bhattacharyya, K. Musier-Forsyth, and S. Hati. 2012. Role of Coupled Dynamics in the Catalytic Activity of Prokaryotic-like Prolyl-tRNA Synthetases. Biochemistry 51:2146–2156.

7. Johnson, J. M., B. L. Sanford, A. M. Strom, S. N. Tadayon, B. P. Lehman, A. M. Zirbes, S. Bhattacharyya, K. Musier-Forsyth, and S. Hati. 2013. Multiple pathways promote dynamical coupling between catalytic domains in Escherichia coli prolyl-tRNA synthetase. Biochemistry 52:4399–4412.

8. Zimmerman, S. B., and S. O. Trach. 1991. Estimation of macromolecule concentrations and excluded volume effects for the cytoplasm of Escherichia coli. J. Mol. Biol. 222:599–620.

9. Ellis, R. J. 2001. Macromolecular crowding: an important but neglected aspect of the intracellular environment. Curr. Opin. Struct. Biol. 11:114–119.

10. Chen, E., A. Christiansen, Q. Wang, M. S. Cheung, D. S. Kliger, and P. Wittung-Stafshede. 2012. Effects of macromolecular crowding on burst phase kinetics of cytochrome c folding. Biochemistry 51:9836–9845.

11. Fan, Y. Q., H. J. Liu, C. Li, Y. S. Luan, J. M. Yang, and Y. L. Wang. 2012. Effects of macromolecular crowding on refolding of recombinant human brain-type creatine kinase. Int. J. Biol. Macromol. 51:113–118.

12. Wang, Y., M. Sarkar, A. E. Smith, A. S. Krois, and G. J. Pielak. 2012. Macromolecular crowding and protein stability. J. Am. Chem. Soc. 134:16614–16618.

13. Dhar, A., A. Samiotakis, S. Ebbinghaus, L. Nienhaus, D. Homouz, M. Gruebele, and M. S. Cheung. 2010. Structure, function, and folding of phosphoglycerate kinase are strongly perturbed by macromolecular crowding. Proc. Nat. Acad. Sci. 107:17586–17591.

14. Batra, J., K. Xu, S. Qin, and H. X. Zhou. 2009. Effect of macromolecular crowding on protein binding stability: modest stabilization and significant biological consequences. Biophys. J. 97:906–911.

15. Qin, S., and H. X. Zhou. 2009. Atomistic modeling of macromolecular crowding predicts modest increases in protein folding and binding stability. Biophys. J. 97:12–19.

16. Pozdnyakova, I., and P. Wittung-Stafshede. 2010. Non-linear effects of macromolecular crowding on enzymatic activity of multi-copper oxidase. Biochim. Biophys. Acta 1804:740–744.

17. Balcells, C., I. Pastor, E. Vilaseca, S. Madurga, M. Cascante, and F. Mas. 2014. Macromolecular crowding effect upon in vitro enzyme kinetics: mixed activation-diffusion control of the oxidation of NADH by pyruvate catalyzed by lactate dehydrogenase. J. Phys. Chem. B 118:4062–4068.

18. Jiang, M., and Z. Guo. 2007. Effects of macromolecular crowding on the intrinsic catalytic efficiency and structure of enterobactin-specific isochorismate synthase. J. Am. Chem. Soc. 129:730–731.

19. Shahid, S., M. I. Hassan, A. Islam, and F. Ahmad. 2017. Size-dependent studies of macromolecular crowding on the thermodynamic stability, structure and functional activity of proteins: in vitro and in silico approaches. Biochim. Biophys. Acta Gen. Subj. 1861:178–197.

20. Kuznetsova, I. M., K. K. Turoverov, and V. N. Uversky. 2014. What Macromolecular Crowding Can Do to a Protein. Int. J. Mol. Sci. 15:23090–23140.

21. Dong, H., S. Qin, and H. X. Zhou. 2010. Effects of macromolecular crowding on protein conformational changes. PLoS Comput. Biol. 6:e1000833.

22. Feig, M., I. Yu, P. H. Wang, G. Nawrocki, and Y. Sugita. 2017. Crowding in Cellular Environments at an Atomistic Level from Computer Simulations. J. Phys. Chem. B 121:8009–8025.

23. Ferreira, L. A., P. P. Madeira, L. Breydo, C. Reichardt, V. N. Uversky, and B. Y. Zaslavsky. 2016. Role of solvent properties of aqueous media in macromolecular crowding effects. J. Biomol. Struct. Dyn. 34:92–103.

24. Burke, B., R. S. Lipman, K. Shiba, K. Musier-Forsyth, and Y. M. Hou. 2001. Divergent adaptation of tRNA recognition by Methanococcus jannaschii prolyl-tRNA synthetase. J. Biol. Chem. 276:20286–20291.

25. Stehlin, C., D. H. Heacock, 2nd, H. Liu, and K. Musier-Forsyth. 1997. Chemical modification and site-directed mutagenesis of the single cysteine in motif 3 of class II Escherichia coli prolyl-tRNA synthetase. Biochemistry 36:2932–2938.

26. Fersht, A. R., J. S. Ashford, C. J. Bruton, R. Jakes, G. L. Koch, and B. S. Hartley. 1975. Active site titration and aminoacyl adenylate binding stoichiometry of aminoacyl-tRNA synthetases. Biochemistry 14:1–4.

27. Heacock, D., C. J. Forsyth, K. Shiba, and K. MusierForsyth. 1996. Synthesis and aminoacyl-tRNA synthetase inhibitory activity of prolyl adenylate analogs. Bioorg. Chem. 24:273–289.

28. Ling, K., H. Y. Jiang, and Q. Q. Zhang. 2013. A colorimetric method for the molecular weight determination of polyethylene glycol using gold nanoparticles. Nanoscale Res. Lett. 8:538.

29. Elstner, M., Q. Cui, P. Munih, E. Kaxiras, T. Frauenheim, and M. Karplus. 2003. Modeling zinc in biomolecules with the self consistent charge-density functional tight binding (SCC-DFTB) method: applications to structural and energetic analysis. J. Comput. Chem. 24:565–581.

30. Frauenheim, T., G. Seifert, M. Elstner, Z. Hajnal, G. Jungnickel, D. Porezag, S. Suhai, and R. Rcholz. 2000. A Self-consistent charge density-functional based tight-binding method for predictive materials simulations in physics, chemistry and biology. Phys. Stat. Sol. B 217:41–62.

31. Berman, H. M., J. Westbrook, Z. Feng, G. Gilliland, T. N. Bhat, H. Weissig, I. N. Shindyalov, and P. E. Bourne. 2000. The Protein Data Bank. Nucl. Acids Res. 28:235–242.

32. Biasini, M., S. Bienert, A. Waterhouse, K. Arnold, G. Studer, T. Schmidt, F. Kiefer, T. Gallo Cassarino, M. Bertoni, L. Bordoli, and T. Schwede. 2014. SWISS-MODEL: modelling protein tertiary and quaternary structure using evolutionary information. Nucl. Acids Res. 42:W252–258.

33. Arnold, K., L. Bordoli, J. Kopp, and T. Schwede. 2006. The SWISS-MODEL workspace: a web-based environment for protein structure homology modelling. Bioinformatics 22:195–201.

34. Benkert, P., M. Biasini, and T. Schwede. 2011. Toward the estimation of the absolute quality of individual protein structure models. Bioinformatics 27:343–350.

35. Humphrey, W., A. Dalke, and K. Schulten. 1996. VMD: visual molecular dynamics. J. Mol. Graph. 14:33–38.

36. Martinez, L., R. Andrade, E. G. Birgin, and J. M. Martinez. 2009. PACKMOL: a package for building initial configurations for molecular dynamics simulations. J. Comput. Chem. 30:2157–2164.

37. Guvench, O., E. Hatcher, R. M. Venable, R. W. Pastor, and A. D. MacKerell. 2009. CHARMM Additive All-Atom Force Field for Glycosidic Linkages between Hexopyranoses. J. Chem. Theor. Comput. 5:2353–2370.

38. Raman, E. P., O. Guvench, and A. D. MacKerell, Jr. 2010. CHARMM additive all-atom force field for glycosidic linkages in carbohydrates involving furanoses. J. Phys. Chem. B 114:12981–12994.

39. Guvench, O., S. S. Mallajosyula, E. P. Raman, E. Hatcher, K. Vanommeslaeghe, T. J. Foster, F. W. Jamison, 2nd, and A. D. Mackerell, Jr. 2011. CHARMM additive all-atom force field for carbohydrate derivatives and its utility in polysaccharide and carbohydrate-protein modeling. J. Chem. Theor. Comput. 7:3162–3180.

40. . Best, R. B., X. Zhu, J. Shim, P. E. M. Lopes, J. Mittal, M. Feig, and A. D. MacKerell. 2012. Optimization of the additive CHARMM all-atom protein force field targeting improved sampling of the backbone φ, ψ and side-chain χ(1) and χ(2) dihedral angles. J. Chem. Theor. Comput. 8:3257–3273.

41. Mallajosyula, S. S., O. Guvench, E. Hatcher, and A. D. Mackerell, Jr. 2012. CHARMM Additive All-Atom Force Field for Phosphate and Sulfate Linked to Carbohydrates. J. Chem. Theor. Comput. 8:759–776.

42. Essmann, U., P. L., and M. L. Berkowitz. 1995. A smooth particle mesh Ewald method. J. Chem. Phys. 103:8577.

43. Glykos, N. M. 2006. Software news and updates. Carma: a molecular dynamics analysis program. J. Comput. Chem. 27:1765–1768.

44. Koukos, P. I., and N. M. Glykos. 2013. Grcarma: A fully automated task-oriented interface for the analysis of molecular dynamics trajectories. J. Comput. Chem. 34:2310–2312.

45. Phillips, J. C., R. Braun, W. Wang, J. Gumbart, E. Tajkhorshid, E. Villa, C. Chipot, R. D. Skeel, L. Kalé, and K. Schulten. 2005. Scalable molecular dynamics with NAMD. J. Comput. Chem. 26:1781–1802.

46. van Aalten, D. M., J. B. Findlay, A. Amadei, and H. J. Berendsen. 1995. Essential dynamics of the cellular retinol-binding protein--evidence for ligand-induced conformational changes. Protein Eng. 8:1129–1135.

47. Brooks, B. R., R. E. Bruccoleri, B. D. Olafson, D. J. States, S. J. Swaminathan, and M. Karplus. 1983. CHARMM: A program for macromolecular energy, minimization, and dynamics calculations. J. Comput. Chem. 4:187–217.

48. Zhang, C. M., J. J. Perona, K. Ryu, C. Francklyn, and Y. M. Hou. 2006. Distinct kinetic mechanisms of the two classes of Aminoacyl-tRNA synthetases. J. Mol. Biol. 361:300–311.

49. Balcells, C., I. Pastor, L. Pitulice, C. Hernández, M. Via, J. L. Garcés, S. Madurga, E. Vilaseca, A. Isvoran, M. Cascante, and F. Mas. 2015. Macromolecular crowding upon in-vivo like enzyme kinetics: Effect of enzyme obstacle size ratio. New Front. Chem. 24:3–16.

50. Wu, J., C. Zhao, W. Lin, R. Hu, Q. Wang, H. Chen, L. Li, S. Chen, and J. Zheng. 2014. Binding characteristics between polyethylene glycol (PEG) and proteins in aqueous solution. J. Mater. Chem. B 2:2983–2992.

51. Lakowicz, J. R. 2006. Protein Fluorescence. Principles of Fluorescence Spectroscopy. J. R. Lakowicz, editor. Springer US, Boston, MA, pp. 529–575.

52. Hixon, J., and Y. Reshetnyak. 2009. Algorithm for the Analysis of Tryptophan Fluorescence Spectra and Their Correlation with Protein Structural Parameters. Algorithms 2:1155.

53. Ghisaidoobe, A. B., and S. J. Chung. 2014. Intrinsic tryptophan fluorescence in the detection and analysis of proteins: a focus on Forster resonance energy transfer techniques. Int. J. Mol. Sci. 15:22518–22538.

54. Lee, J. C., and S. N. Timasheff. 1981. The stabilization of proteins by sucrose. J. Biol. Chem. 256:7193–7201.

55. Yadav, R., and P. Sen. 2013. Mechanistic investigation of domain specific unfolding of human serum albumin and the effect of sucrose. Protein Sci. 22:1571–1581.

56. Paul, S. S., P. Sil, R. Chakraborty, S. Haldar, and K. Chattopadhyay. 2016. Molecular Crowding Affects the Conformational Fluctuations, Peroxidase Activity, and Folding Landscape of Yeast Cytochrome c. Biochemistry 55:2332–2343.

57. Laskowski, R. A., M. W. MacArthur, D. S. Moss, and J. M. Thornton. 1993. PROCHECK: a program to check the stereochemical quality of protein structures. J. Appl. Crystallogr. 26:283.

58. Laskowski, R. A., J. A. Rullmannn, M. W. MacArthur, R. Kaptein, and J. M. Thornton. 1996. AQUA and PROCHECK-NMR: programs for checking the quality of protein structures solved by NMR. J. Biomol. NMR 8:477–486.

59. Wimmerova, M., S. Kozmon, I. Necasova, S. K. Mishra, J. Komarek, and J. Koca. 2012. Stacking interactions between carbohydrate and protein quantified by combination of theoretical and experimental methods. PLoS One 7:e46032.

60. Spiwok, V. 2017. CH/pi Interactions in Carbohydrate Recognition. Molecules 22.

61. Kim, S., S. W. Lee, E. C. Choi, and S. Y. Choi. 2003. Aminoacyl-tRNA synthetases and their inhibitors as a novel family of antibiotics. Appl. Microbiol. Biotechnol. 61:278–288.

62. Lv, P. C., and H. L. Zhu. 2012. Aminoacyl-tRNA synthetase inhibitors as potent antibacterials. Curr. Med. Chem. 19:3550–3563.

63. Dewan, V., J. Reader, and K. M. Forsyth. 2014. Role of aminoacyl-tRNA synthetases in infectious diseases and targets for therapeutic development. Top. Curr. Chem. 344:293–329.

64. Pham, J. S., K. L. Dawson, K. E. Jackson, E. E. Lim, C. F. Pasaje, K. E. Turner, and S. A. Ralph. 2014. Aminoacyl-tRNA synthetases as drug targets in eukaryotic parasites. Int. J. Parasitol. Drugs Drug Resist. 4:1–13.

65. Schultz, S. G., and A. K. Solomon. 1961. Determination of the effective hydrodynamic radii of small molecules by viscometry. J. Gen. Physiol. 44:1189–1199.

